# Optimizing the identification of causal variants across varying genetic architectures in crops

**DOI:** 10.1101/310391

**Authors:** Chenyong Miao, Jinliang Yang, James C. Schnable

## Abstract

**Background:** Association studies use statistical links between genetic markers and variation in a phenotype’s value across many individuals to identify genes controlling variation in the target phenotype. However, this approach, particularly conducted on a genome-wide scale (GWAS), has limited power to identify the genes responsible for variation in traits controlled by complex genetic architectures.

**Results:** Here we employ simulation studies utilizing real-world genotype datasets from association populations in four species with distinct minor allele frequency distributions, population structures, and patterns linkage disequilibrium to evaluate the impact of variation in both heritability and trait complexity on both conventional mixed linear model based GWAS and two new approaches specifically developed for complex traits. Mixed linear model based GWAS rapidly losses power for more complex traits. FarmCPU, a method based on multi-locus mixed linear models, provides the greatest statistical power for moderately complex traits. A Bayesian approach adopted from genomic prediction provides the greatest statistical power to identify causal genetic loci for extremely complex traits.

**Conclusions:** Using estimates of the complexity of the genetic architecture of target traits can guide the selection of appropriate statistical methods and improve the overall accuracy and power of GWAS.

## Background

Association studies have been widely adopted as a complement forward and reverse genetics approaches in identifying and characterizing the functions of specific genes. Unlike forward and reverse genetics approaches, association studies also identify functionally variable alleles segregating in target populations. These alleles can guide breeding efforts in crop and livestock species, as well as providing increasingly accurate predictions of disease risk factors in humans. Advances in genotyping technology have dramatically reduced the barriers to conducting association studies with genome-wide genetic marker datasets across natural populations. Since becoming feasible mid-2000s, Genome Wide Association Studies (GWAS) have been successfully used to identify thousands of single nucleotide polymorphisms (SNPs) associated with diseases in human [1] and complex agricultural traits in plants [2, 3, 4, 5]. For most traits analyzed, loci identified by GWAS can generally explain only a subset of total genetically controlled phenotypic variation for most traits analyzed [6, 7, 8]. Many explanations have been proposed for this “missing heritability” including epigenetic effects [9], epistasis [10, 11, 12], structural variants which are not detected by conventional SNP genotyping [13], rare alleles with large effects, and common alleles small effect sizes [14, 15]. While the first two proposed explanations for missing heritability are more difficult to address, both rare alleles with large effect sizes and common alleles with small effect sizes can potentially be identified through increases in the statistical power of GWAS to identify causal variants.

Many traits of interest to biologists are controlled by complex genetic architectures [3, 5, 16] where hundreds, thousands, or the majority of all genes [17] may control variation in the target trait. The most straightforward approach to increasing the proportion of causal identified is to increase the size of genotyped and phenotyped populations. However, increases in population size are expensive, and subject to diminishing returns in terms of the improvement of power to detect both rare alleles and alleles with small effect sizes. Improved statistical approaches to isolating a larger proportion of total causal variants controlling complex traits is therefore highly desirable.

Currently GWAS approaches based on mixed linear models (MLM) are widely employed in both plant and animal systems. MLM based models are able to control for confounding effects of both population structure and unequal relatedness between individuals which are left uncontrolled in approaches based on Generalized Linear Models (GLM), at the expense of greater run times. A wide range of different algorithms have been proposed and developed to improve the computational efficiency of MLM models, including EMMAX [18], Compressed-MLM [19], FaST-LMM [20], and GEMMA-MLM [21]. However, because MLM-based methods are ultimately evaluating the relationship between each genetic marker and the overall variation in a given trait across a population independently, the statistical power of these methods rapidly decreases as the total number of genes controlling variation in a given trait increases, and the proportion of total genetic variance explained by any one locus decreases.

Multi-locus mixed-models (MLMM) explicitly identify and control for the effects of large effect loci as fixed effects as these loci are identified by the model [22]. This approach increases the proportion of the remaining genetic variance explained by the remaining unidentified loci, and increases the statistical power of the method to detect a greater number of causal variants for complex traits. While the high computational cost of MLMM initially acted as a barrier to widespread adoption, a modified MLMM method, Fixed and random model Circulating Probability Unification (FarmCPU), has dramatically reduced the computational complexity and computing time of this approach [23], and ongoing optimization and parallelization efforts have continued to decrease real-world run times for MLMM based approaches [24].

A second potential approach to accurately identifying individual causal variants for traits controlled by complex genetic architectures is the use of Bayesian multiple-regression methods. The Bayesian-based models fit all the available markers simultaneously, which makes them especially suitable to study highly polygenic traits. While Bayesian approaches including BayesA, BayesB, BayesC, and BayesCπ has been widely employed in genomic prediction and selection [25, 26, 27, 28], in principal the same statistical approaches can be employed to identify the individual loci controlling variation in target traits [29]. Several studies have employed Bayesian multiple-regression methods to identify putative causal variants [30, 31, 32], however the performance of these bayesian methods when employed in GWAS have not been extensively evaluated relative to current non-bayesian approaches.

Here we compare the performance of MLM, MLMM (FarmCPU), and Bayesian GWAS approaches across simulated trait datasets containing 2 - 1024 causal variants and heritability ranging from 0.1 to 1. To capture realistic patterns of minor allele frequency distributions, population structure and linkage disequilibrium, real world genotype datasets from four widely studied crop species: rice (*Oryza sativa*), foxtail millet (*Setaria italica*), sorghum (*Sorghum bicolor*), and maize (*Zea mays*) [4, 5, 33, 34] (Table 1). We demonstrate that the power and accuracy of both FarmCPU and BayesCπ to identify causal variants for complex traits exceed conventional MLM approaches. Of the three methods, FarmCPU generally provides the most favorable trade off between power and low false positive rates for moderately complex traits controlled by several dozen loci, while the BayesCπ approach provides a more favorable trade off for traits controlled by hundreds of loci. However, the number of casual loci where the crossover between the comparative advantages of these two methods occurred varied across species. The results presented here, including a set of 4,000 simulated phenotypic datasets generated using real world genotype datasets, will provide both a resource for evaluating future innovations in GWAS software, and information to help researchers select the most effective experimental design and statistical approach for their particular research projects.

**Table 1.** Statistical summary of each genotype dataset.

| Species | Genotyping technology | Genome size (Mb) | LD Decay (Kb) | Number of Accessions | Number of Markers |
| --- | --- | --- | --- | --- | --- |
| *Sorghum bicolor* | GBS | 732 | 2 | 2,327 | 354,940 |
| *Setaria italica* | Low coverage WGS | 406 | 794 | 916 | 663,985 |
| *Oryza sativa* | Microarray | 372 | 0.004 | 1,568 | 629,019 |
| *Zea mays* | GBS | 2,300 | 0.063 | 2,503 | 560,515 |

## Results

Each of the four populations employed in this study presented a different combination of linkage disequilibrium, minor allele frequency distribution, and population structure (Figure 1, Table 1). These differences may result from differences in population demographics, criteria used to assemble the populations, and genotyping technologies employed each of the genotype datasets. For example, the comparatively low frequency of rare alleles in rice results from selection loci with more frequent minor alleles prior to microarray design [34], while the low frequency of rare alleles in foxtail millet resulted from a post-genotyping, prepublication filter for loci with relatively more common minor alleles [4]. With the exception of rice, where an absence of strong LD was incorporated into the selection criteria for markers prior to genotyping [34], the pattern of LD decay observed across populations of the remaining three species exhibited a negative correlation with reported outcrossing frequencies for each species, suggesting that this difference is the result of biological variation rather than genotyping strategy [35, 36, 37, 38, 39] (Figure S1).

**Figure 1.**
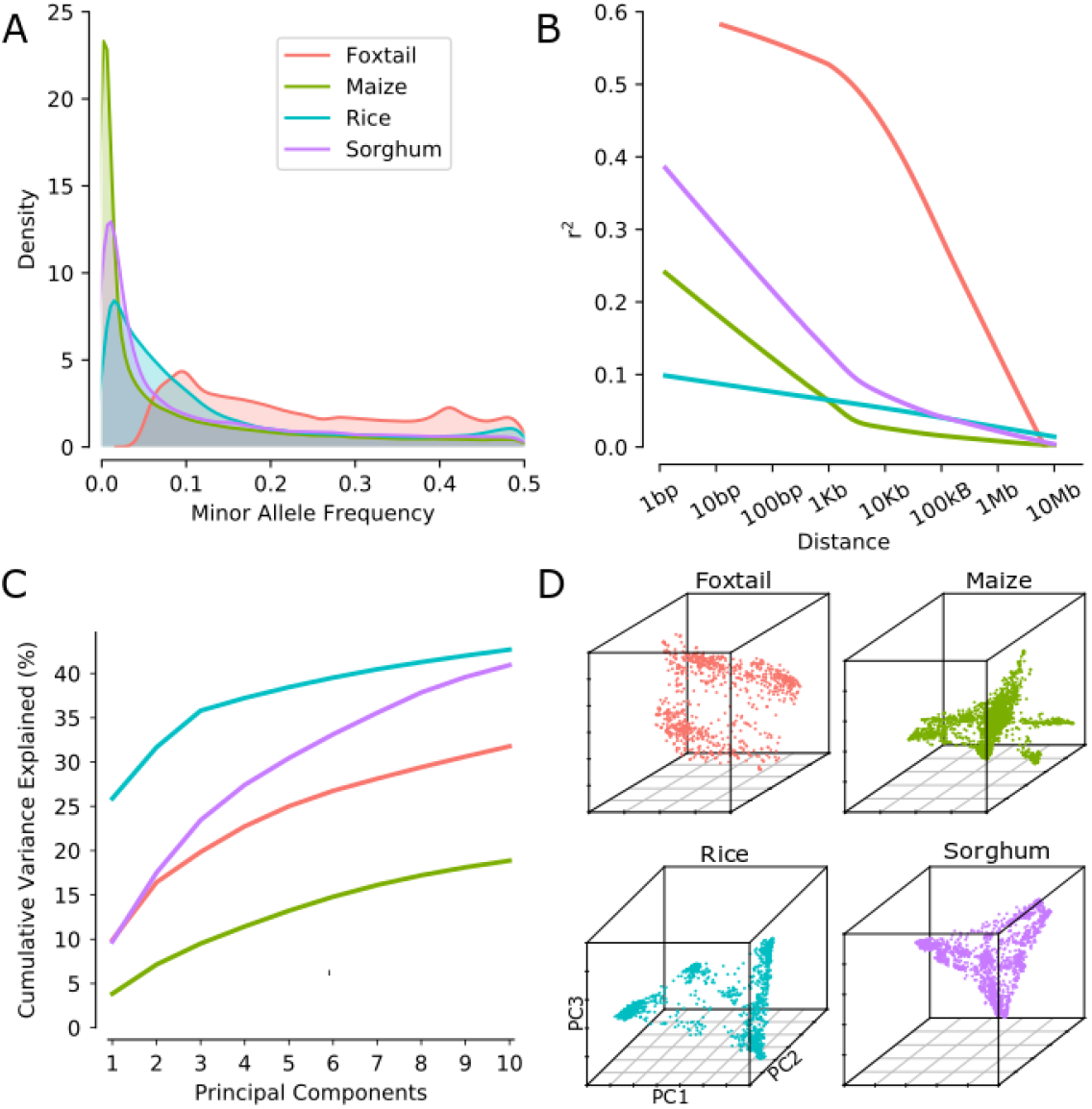
Characterization of the four association populations and associated genotype datasets employed in this study. (A) Distribution of minor allele frequencies across all genotyped markers in each population. (B) Patterns of linkage disequilibrium decay in each population based on average pairwise *r*^2^ between genetic markers (Methods). (C) Cumulative proportion of total genotypic variance explained up to ten principal components in each population. (D) PCA distribution for individuals in each population.

### Evaluation mixed linear model based GWAS

A total of 1,000 phenotype datasets were generated per species, with ten independent replicates with different sets of causal SNPs for each possible combination of 10 different levels of heritability and 10 different levels genetic architecture complexity. A causal variant was considered to be identified if either the causal genetic marker selected by the simulations, or one or more markers with an LD ≥ with the causal SNP was identified by a given GWAS analysis. As expected, power to detect true positives decreased in response to both increases in the number of simulated causal variants controlling the trait and decreases in simulated heritability. The MLM-based approach failed to identify the vast majority of causal variants for traits controlled by 256 or more loci (Figure 2). Consist with both theory and previous studies both rare alleles and alleles assigned smaller absolute effect sizes were the least likely to be identified in the MLM based GWAS analysis (Figure S4, Table S1). Subsampling of each population was used to evaluation how rapidly the proportion of total causal SNPs identified increases with increased population size. The effect of increasing population size was relatively more pronounced when genetic architecture was less complex, and smaller increases were observed with increasing population size for more complex genetic architectures controlled by >100 causal variants (Figure 3, Figure S5).

**Figure 2.**
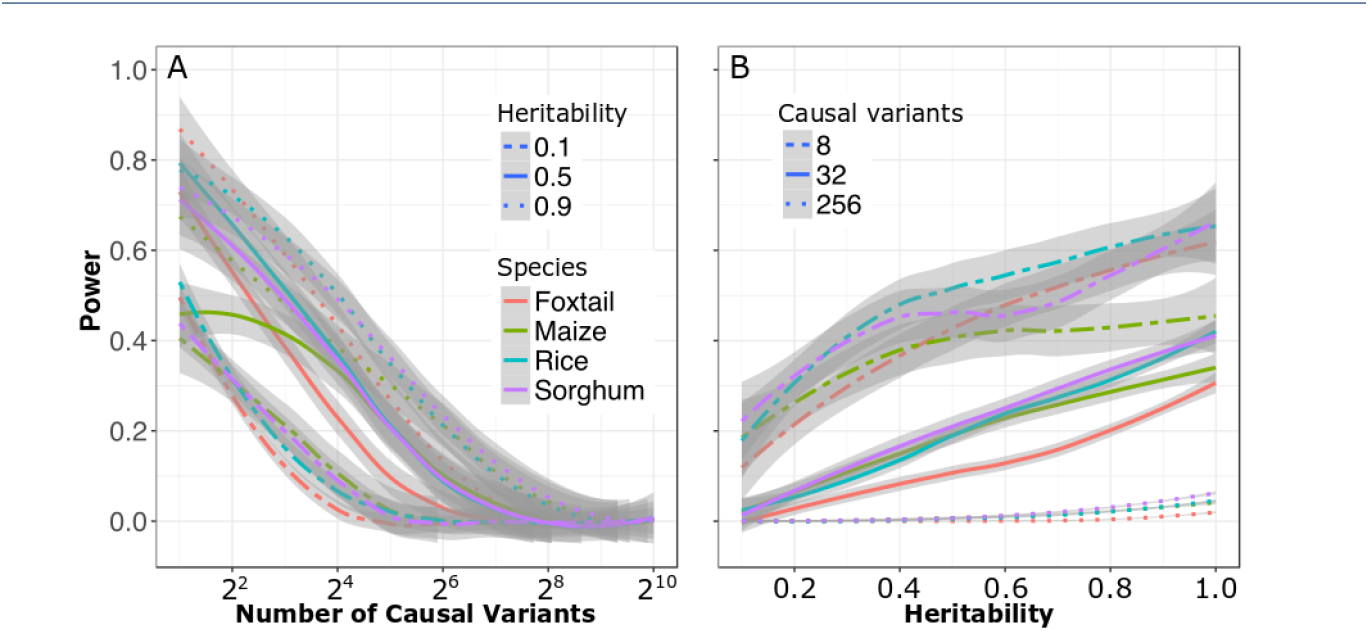
Changes in the power of conventional (MLM based) GWAS to identify causal variants in response to changes in heritability and the complexity of the genetic architecture controlling the target trait. Data shown is from foxtail millet. Comprehensive results from all four populations in Supplementary Figures S2, S3. (**A**) Change in power to detect true positives as the number of causal variants increases under high (0.9), medium (0.5), and low (0.1) levels of heritability. (**B**) Change in power to detect true positives as heritability decreases for traits controlled by simple (N=8), moderately complex (N=32), and complex (N=256) genetic architectures. Positive calls were defined as those above a Bonferroni corrected p-value cutoff of 0.05.

**Figure 3.**
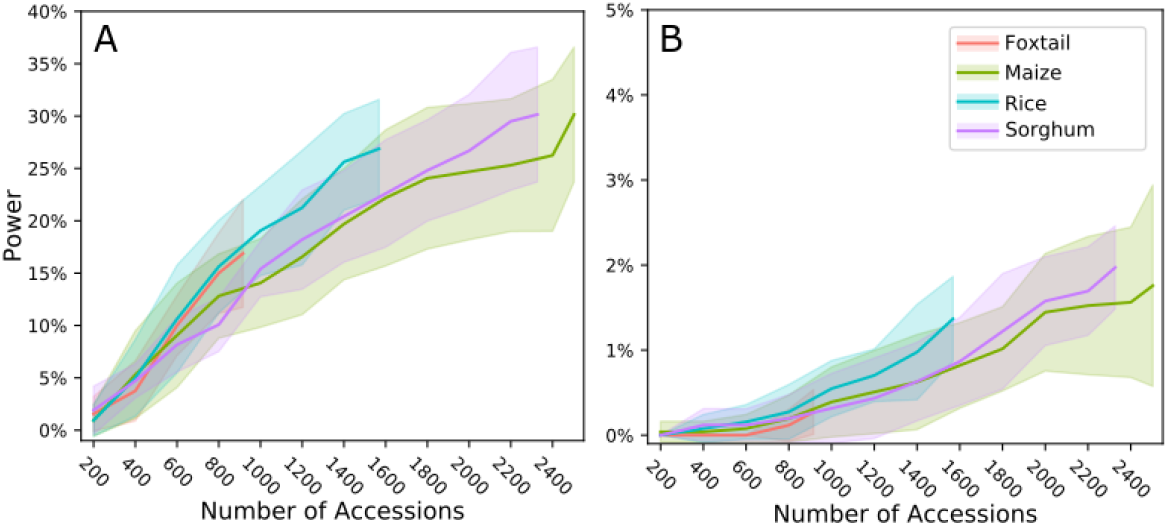
Changes in the power of conventional (MLM-based) GWAS to identify causal variants for complex traits in response to increases in population size in each of the four association populations evaluated. (A) a moderately complex trait controlled by 32 loci (B) a complex trait controlled by 256 loci. Both analyses used data from traits with heritability of 0.7. Positive calls were defined as those above a Bonferroni corrected p-value cutoff of 0.05.

### Alternative GWAS methods for complex traits

As shown above, MLM-based GWAS identifies only a small proportion of causal genetic variants for complex traits controlled by hundreds or thousands of distinct genetic loci. We next evaluated two methods specifically developed to analyze polygenic trait: FarmCPU [23] and BayesCπ [40]. To avoid confounding effects from different approaches to scoring the strength of associations between markers and trait variation, cross-method comparisons are made based on selecting equivalent numbers of positive genetic markers in each analysis. The proportion of causal variants detected for a given total number of positive genetic markers declines in each species as heritability decreases and as the total number of causal variants controlling the trait increases. However, FarmCPU and BayesCπ both consistently outperformed MLM-based analysis in terms of both overall proportion of causal variants identified and FDR control (Figure 4, 5). For moderately complex traits (QTN = 32, 64) the statistical power of BayesCπ and FarmCPU provided approximately equivalent statistical power, however FarmCPU tends to provide lower false discovery rates than BayesCπ for these genetic architectures. For complex traits (128, 256 causal variants), the BayesCπ approach outperforms FarmCPU on both power and false discovery rate metrics (Figures 4, 5, S7, S9), although the difference in performance between the two methods is less apparent for traits with lower levels of heritability (Figures S10, S11, S12, S13)

**Figure 4.**
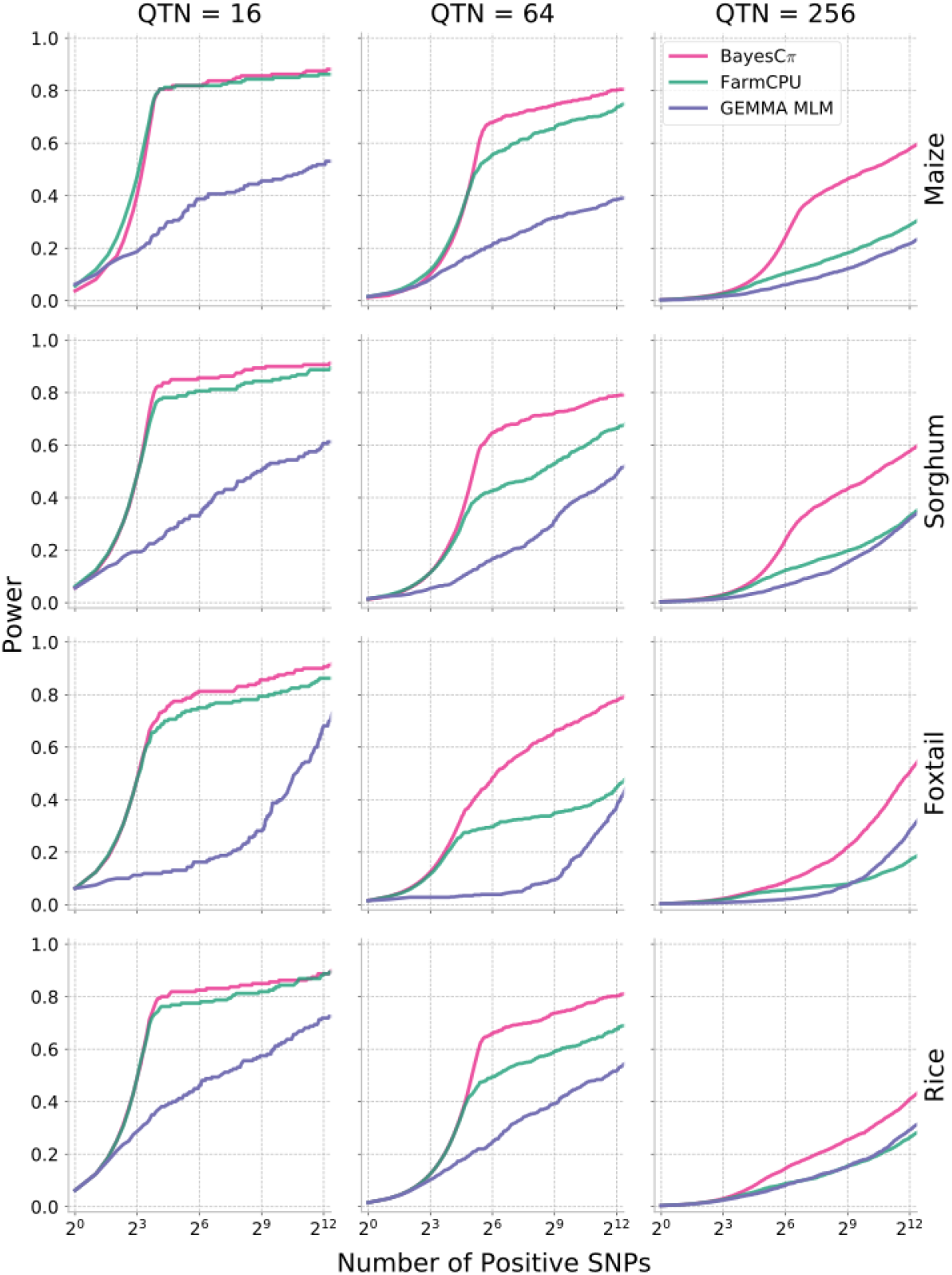
Changes in the power of three statistical approaches to GWAS across all four association populations in response to changes in changes in the statistical threshold employed. To enable comparisons across different methods with different approaches to reporting statistical significance, the x-axis is ordered by the total number of positive genetic markers accepted at a given statistical threshold. Data shown is for traits with increasingly complex genetic architectures with near-best-case assumptions for trait heritability (0.9). Results for all other simulated genetic architectures are provided in Supplemental Figures S6, S7.

**Figure 5.**
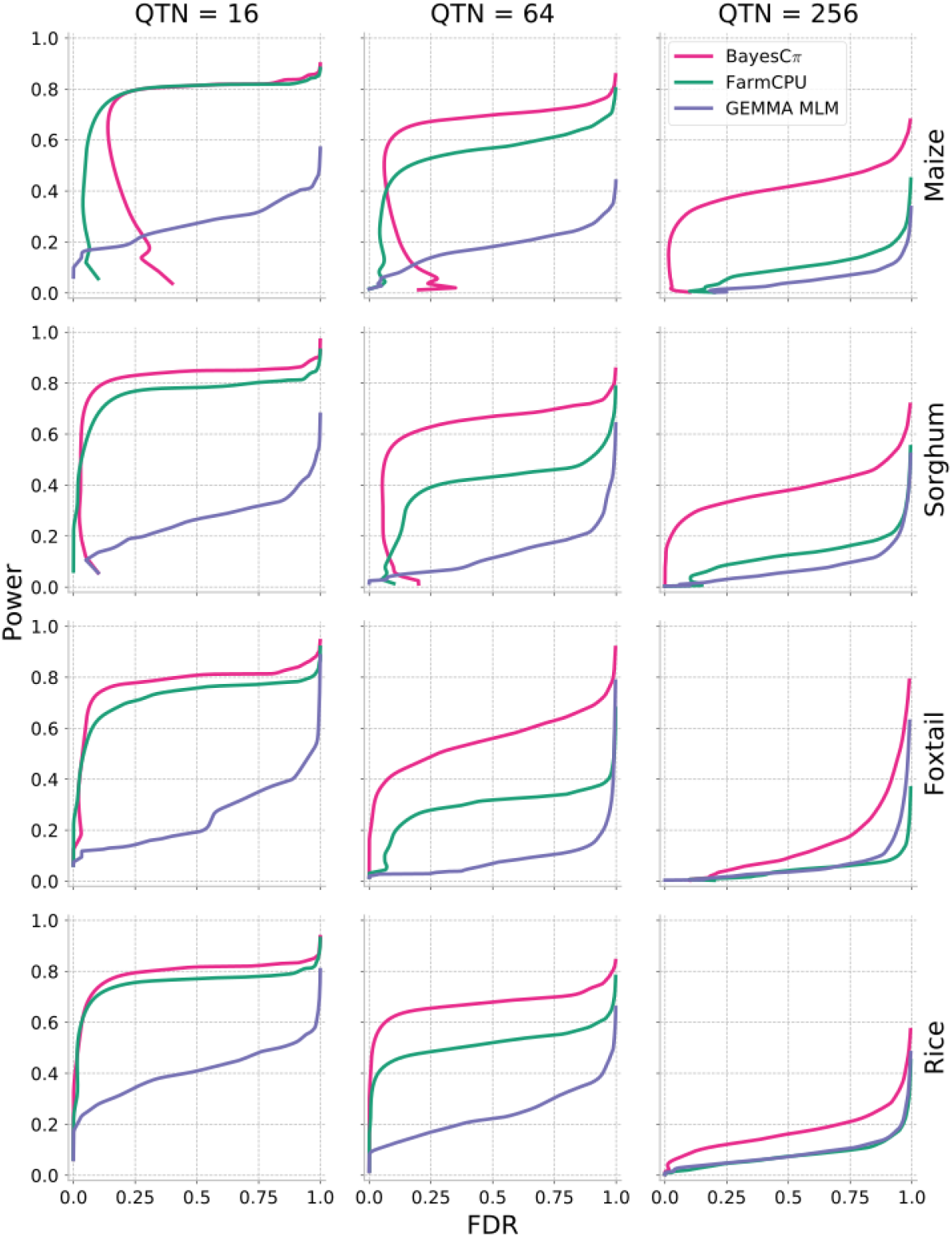
Relationship between power and false discovery rate using each GWAS method to analyze simple, medium and complex traits in each population. Data shown is for traits with increasingly complex genetic architectures with near-best-case assumptions for trait heritability (0.9). Results for all other simulated genetic architectures are provided in Supplemental Figures S8, S9.

For sets of simulation parameters where BayesCπ and FarmCPU identified equivalent proportions of true causal variants, the specific causal variants identified by each algorithm were different. Causal SNPs classified into four mutually exclusive categories: those identified by both algorithms, those identified by either only FarmCPU or only BayesCπ, and those missed by both. Causal SNPs identified by both algorithms tended to have higher MAFs (p=0.0197, p=0.0065, shared vs FarmCPU only, shared vs Bayesian only, Mann-Whitney U test), and larger effect sizes (p=1.17e-05, p=1.23e-14, shared vs FarmCPU only, shared vs Bayesian only, Mann-Whitney U test). SNPs identified only by FarmCPU tended to have lower MAF frequencies than those identified only by Bayesπ (p=0.0008, Mann-Whitney U test), and did not exhibit a statistically significant different in effect size (p=0.2185, Mann-Whitney U test). Overall, the two approaches appear to have complementary strengths for identifying different subsets of allelic variants missed by conventional GWAS methods.

**Figure 6.**
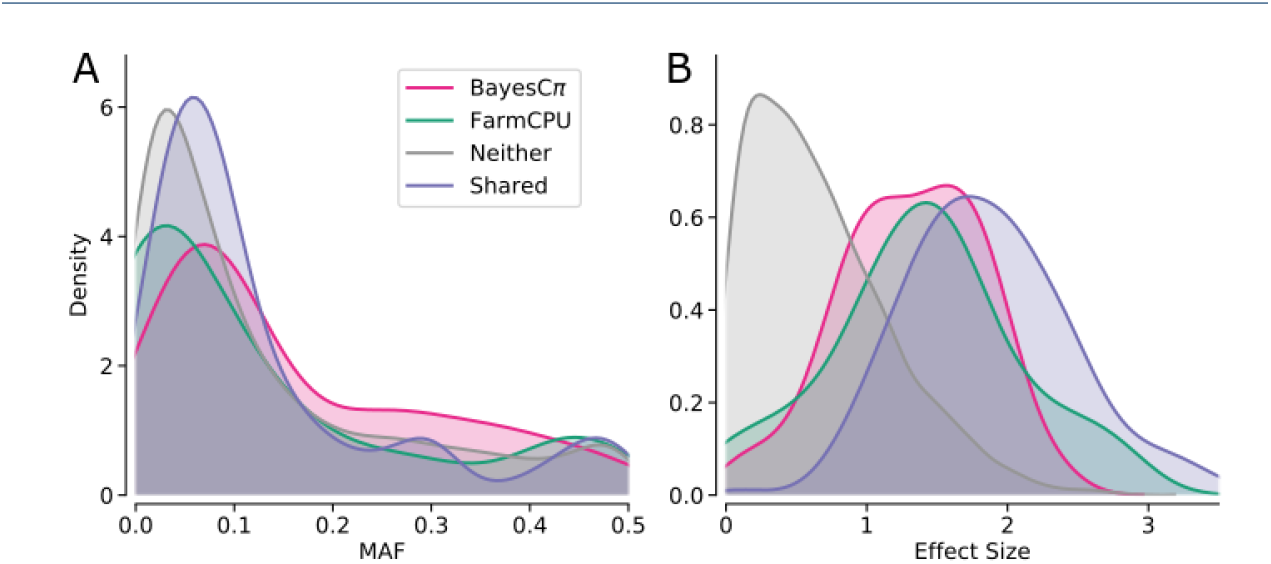
Differences in the characteristics of causal SNPs identified using either BayesCπ or FarmCPU. Distribution of MAF (A) and absolute effect size (B) for causal variants identified by both BayesCπ or FarmCPU, only Bayesπ, only FarmCPU, or neither method. Data shown are from the rice population, with 256 total causal variants per simulation, and a heritability of 0.7. Equivalent analyses from other species and varying levels of heritability are provided in Supplemental Figure S14 and Supplemental Table S2.

## Discussion

In this study, we employed four genotype datasets with different population structures from different crop species. Mixed linear models showed substantial reductions in power as the complexity of the genetic architecture of the trait being analyzed increased. Two approaches developed to address the challenge of GWAS in traits controlled by complex architectures (FarmCPU, a modified MLMM) and an approach based on BayesCπ show complementary strengths and higher power and lower false discovery rates for complex traits. FarmCPU provided a more favorable trade off between power and FDR for moderately complex traits and a greater likelihood of identifying rare causal variants, while BayesCπ based GWAS provided greater power for extremely complex traits and a greater likelihood of identifying variants with smaller absolute effect sizes but which are more common within the study population. Present statistical approaches to GWAS have the greatest statistical power to identify SNPs which are both common, and control a large proportion of total genetic variation in the target populations. As a result, few previously unknown loci with utility for plant breeding have been discovered through GWAS-based analysis [41]. The identification of common alleles with moderate effect sizes and rare alleles with large effects would improve the utility for GWAS for both basic biological and applied applications.

Methods to determine the complexity of the genetic architectures controlling different traits are clearly also needed, given the differences in the relative strengths of MLM, MLMM and Bayesian approaches for traits controlled by genetic architectures different levels of complexity. The BayesCπ method includes a statistical approach to estimate the number of causal variants controlling a given trait prior to fitting a model to the data [40]. These estimates serve as a prior for model fitting in BayesCπ. However, as different GWAS approaches provide the most favorable results for traits with different complexities, estimation of the number of genetic loci controlling a trait can also guide which statistical approach is best suited to analyze a given dataset. The accuracy of the estimates of the number of causal variants generated by the BayesCπ algorithm were estimated across varying levels of heritability and trait complexity. In all four species, while BayesCπ was able to accurately estimate heritability for traits controlled by different numbers of causal variants S15 and also provided relatively accurate and unbiased estimated of the number of causal variants when the heritability of the trait was high and/or the total number of causal variants was smaller, the number of causal variants was systematically overestimated for complex traits with lower levels of heritability (Figure 7). The reduced accuracy in estimating the number of causal variants for traits with low levels of heritability and complex genetic architectures may explain why the differences in performance between FarmCPU and BayesCπ is less apparent analyses of datasets will lower heritability. One potential explanation for this observation is that the model is attempting to explain residual error - not heritable phenotypic variation – by including additional, noncausal SNPs in the model. However, with awareness of this limitation, estimation of the number of causal variants controlling a given trait can aid researchers in determining which GWAS method is likely to provide the most informative result for a given dataset.

**Figure 7.**
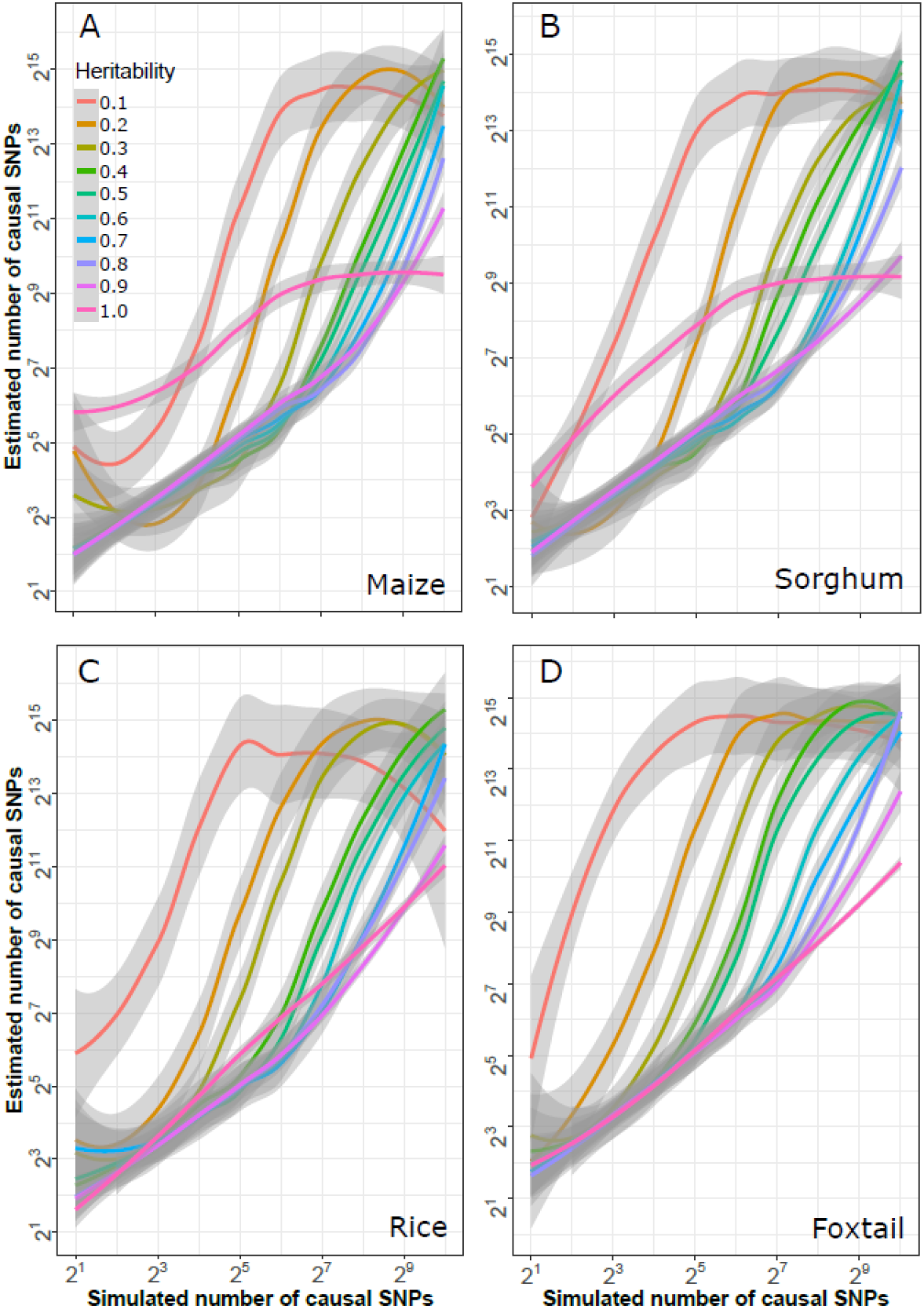
Relationship between estimated complexity of genetic architectures generated by BayesCπ and true genetic architecture complexity given different levels of heritability in each species. Grey areas indicate 95% confidence bands around each estimate.

Evaluations of GWAS approaches can be performed using either real data or simulated data. Here simulated data was employed, as it provided comprehensive ground truth information for comparison across methods, something unavailable for real world phenotype datasets for complex traits. The use of real world genotype datasets captured minor allele frequency distributions and population structures comparable to those observed in the real world. However, it is also important to acknowledge the limitations of simulation based studies. The simulated phenotype datasets employed here assumed the effect sizes of minor alleles are drawn from a normal distribution, which is supported by real world observations (as shown in Figure S18) [42, 43]. More significantly, the simulation parameters used assumed no correlation between the minor allele frequency of an allele and its effect size, which does not match predictions from population genetic theory or observation that rare alleles tend to be associated with larger molecular phenotypes in maize [44] In addition, the statistical model used to generate phenotype datasets here did not incorporate epistatic interactions between causal variants. Finally, we did not model errors in genotyping or phenotyping, and thus cannot evaluate whether the different statistical approaches to GWAS evaluated here show different levels of robustness to errors in data collection.

While the results presented here for the use of BayesCπ to identify causal variants are promising, additional work is needed to further adapt BayesCπ for use in GWAS applications. The model employed here did not yet incorporate any controls for population structure. Integrating such a control might marginally reduce power, but should substantially reduce type I error rates in the Bayesian model. The model we employed also provided a ranking of genetic markers but not a straightforward method of establishing a cutoff between candidate causal variants and non-candidate loci. While ranking enabled comparisons of power, type I error, and false discovery rate, the application of BayesCπ-based GWAS in a real world setting will require methods to establish such cutoffs. One promising approach recently discussed in the literature is to estimate posterior type I error rates [45]. Approaches using machine learning to identify cutoffs, such in NeuralFDR, also seem a promising avenue of investigation [46]. In addition computational resource requirements play an substantial role in which statistical approaches become widely adopted over time. With the largest of the four genotype datasets employed here (maize) BayesCπ required approximately 4.5 Gb of RAM and 2 hours to analyze one dataset. For comparison, the MLM implementation in GEMMA required only 1 Gb of RAM and approximately 40 minutes to analyze the same dataset and FarmCPU required approximately 30 minutes and 5.5 Gb RAM (Supplemental Figures S16, S17). However, optimization of computational pipelines can reduce run times dramatically without the need for changes to statistical models. Modifications to the reference implementation of the FarmCPU algorithm have been shown to produce the same results while reducing runtime by approximately two-thirds [24].

## Conclusion

Association studies have been and seem likely to remain an important tool for investigating how genotype determines phenotype. While certain diseases and target traits for breeding efforts are controlling by a small number of large effect loci segregating in Mendelian fashion, many traits of interest are controlled by moderately or extremely complex genetic architectures. Here we have shown that different approaches to GWAS have complementary strengths, and the complexity of the genetic architecture controlling a target trait should be determined prior to the selection of an appropriate statistical approach for analyzing a given dataset. Further improvements in both statistical approaches and computational optimization hold the promise of dramatically expanding our understanding of the role that both rare alleles with large consequences and common alleles with small consequences play in determining how genotype determines phenotype across species.

## Methods

### Genotype dataset sources and filtering parameters

Genotype dataset for foxtail millet (*Setaria italica*) [4], maize (*Zea mays*) [33], sorghum (*Sorghum bicolor*)[5], and rice (*Oryza sativa*) [34] were taken from published sources. Foxtail millet SNPs were discovered and scored using low coverage (0.5x) whole-genome resequencing reads aligned to the *Setaria italica* reference genome (v2 from Phytozome v7.0) [47]. The partially imputed SNP dataset was downloaded from Millet GWAS Project website: http://202.127.18.221/MilletHapl/GWAS.php [4]. The downloaded genotype data included 916 diverse varieties and 726,080 SNPs. SNPs with minor allele frequencies lower than 5% had been removed prior to the publication of the dataset. After downloading, SNPs without calls in >10% of samples were removed from the dataset. A sorghum GBS dataset which included 404,627 SNPs scored relative to the v1.4 of the sorghum reference genome [48] across a set of 1,943 accessions were downloaded from Data Dryad http://datadryad.Org/resource/doi:10.5061/dryad.jc3ht/1 [5].After downloading, SNPs without genotype calls in >30% of samples and SNPs with heterozygous calls in > 5% of samples with genotype calls were removed from the dataset. A maize GBS dataset which included calls for 681,257 SNPs relative to B73_RefGen_V1 [49] across a set of 2,815 accessions was downloaded from Panzea https://www.panzea.org/ [33]. After downloading, SNPs without genotype calls in >30% of samples and SNPs with heterozygous calls in >5% of samples with genotype calls were removed from the dataset. After any filtering parameters described above for individual datasets, missing data points in foxtail millet, sorghum, and maize dataset were imputed using Beagle v4.1 with default parameters [50]. Data from genotyping 1,568 diverse rice accessions using the 700,000 marker HDRA microarray platform was downloaded from GEO (ID: GSE71553) [34]. After download SNPs with heterozygous genotype calls in >5% of samples were removed from the dataset. Statistics on the final number of marker and samples in each dataset are provided in Table 1.

### Characteristics and summary statistics of genotype datasets used in this study

The minor allele frequency was calculated for each SNP in each dataset. Patterns of minor allele frequency distributions for each dataset were assessed and visualized using kernel density plots generated using the the function ‘kdeplot’ from the Python package ‘seaborn.’ For each dataset, the top ten principal components were calculated using Tassel 5.2 [51]. The top three principal components from the same analyses were used to plot population structure using the R package scatterplot3d. Plink 1.9 was used to calculated *r*^2^ between all pairs of SNP markers separated by less than 10 kilobases [52]. For windows greater than 10 kilobases markers were subsampled to retain approximately equal numbers of pairwise comparisons in the 0-10 kilobase internal, in the 10 to 100 kilobase internal, in the 100 kilobases to 1 megabase interval, and in the 1 megabase to 10 megabase internal. In each dataset average *r*^2^ values were calculated from 10^0 1^ to 10^7^ using a logarithmic step size of 0.1. A regression curve was fit to these values using the function ‘regplot’ from the Python package ‘seaborn.’

### Phenotype simulation

Phenotype datasets were simulated using an additive genetic model derived from the underlying genotype datasets:

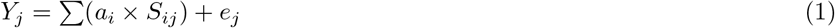

In the model, *Y_j_* is the simulated phenotype for plant *j*; *a_i_* is the effect of the *i*th causal SNP; *S_ij_* is the SNP genotype (coded with 0, 1, 2) for the *i*th causal SNP of the *j*th plant; and *e_j_* is the normal residual error for *j*th plant with mean of 0 and standard deviation of 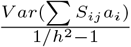, where *h*^2^ denotes the heritability.

An R function ‘simcrop’ was implemented within the open source ‘g3tools’ R package (https://github.com/jyanglab/g3tools). Results presented above employ phenotype datasets where the effect value for each causal SNP was drawn from a normal distribution, however the software package also provides the option to specify other effect size distribution models. Phenotype datasets were simulated for scenarios where the number of causal genetic loci ranged from 2^1^ to 2^10^ (2 ~ 1,024 QTNs) and where the heritability of trait values ranged 0.1 to 1.0 in steps of 0.1. For each combination of heritability and number of causal variants, ten independent datasets with different randomly selected causal variants were selected.

### Genome-wide association studies

The kinship matrix applied in all methods was generated using “-gk 1” option in GEMMA for each genotype dataset. All GEMMA-MLM based GWAS analysis were performed using the “gemma” command (version 0.95alpha) with options “-lmm” and “-k” to specify kinship matrix [21]. FarmCPU was run using the command FarmCPU(Y=myY, GD=myGD, GM=myGM, CV=myCV, method.bin=“optimum”) in R. The parameter method.bin=“optimum” allows the FarmCPU to selected optimized possible QTN window size and number of possible QTNs in the model [23]. The BayesCπ was conducted using GenSel software package [40]. For each run, chain length was set to 11,000 with the first 1,000 as burn in, and π=0.9999. All the GWAS jobs were run on HCC’s (the Holland Computing Center) Crane cluster at University of Nebraska-Lincoln.

## List of abbreviations

GWAS: : Genome-Wide Association Study
GBS: : Genotyping-By-Sequencing
PCA: : Principal Component Analysis
LD: : Linkage Disequilibrium
SNP: : Single Nucleotide Polymorphism
MAF: : Minor Allele Frequency
QTN: : Quantitative Trait Nucleotide
GEMMA: : Genomic Association and Prediction Integrated Tool
GLM: : General Linear Model
MLM: : Mixed Linear Model
MLMM: : Multi-Locus Mixed-Model
FDR: : False Discovery Rate
HDRA: : High-Density Rice Array
HCC: : the Holland Computing Center

## Declarations

### Availability of data and materials

All simulated phenotypes and the corresponding information of causal SNPs are available on figshare http://doi.org/10.6084/m9.figshare.6026246.

### Author’s contributions

JS designed the project. JY generated phenotype datasets and conducted the BayesCπ analysis. CM conducted all other analysis. CM, JY, and JS generated figures and wrote the manuscript. All authors have read and approved the final manuscript.

## Acknowledgements

This material is based upon work supported by the National Science Foundation under Grant No. OIA-1557417 to JCS, award #2016-67013-24613 and work supported by the National Institute of Food and Agriculture, U.S. Department of Agriculture under Award 16-67013-24613 to JCS, and startup funds and a Layman seed award from the University of Nebraska-Lincoln to JY. This work was completed utilizing the Holland Computing Center of the University of Nebraska, which receives support from the Nebraska Research Initiative.

---

**Table S1.**
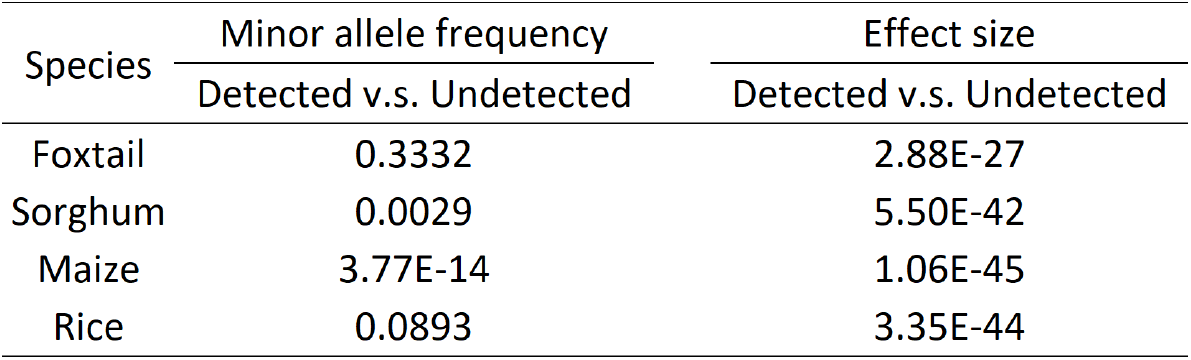
Mann-Whitney U test between SNP groups detected and undetected by GEMMA. Data shown are from simulations where the number of causal variants is 64 and heritability is 0.7

| Species | Minor allele frequency | Effect size |
| --- | --- | --- |
|  | Detected v.s. Undetected | Detected v.s. Undetected |
| Foxtail | 0.3332 | 2.88E-27 |
| Sorghum | 0.0029 | 5.50E-42 |
| Maize | 3.77E-14 | 1.06E-45 |
| Rice | 0.0893 | 3.35E-44 |

**Table S2.**
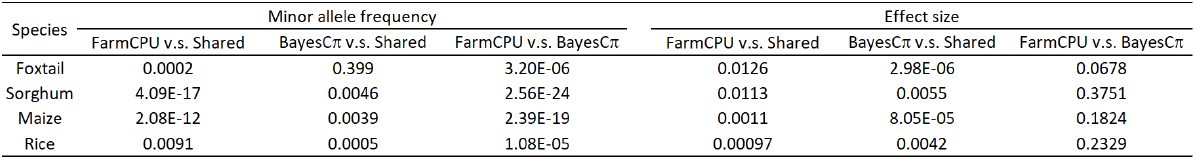
Mann-Whitney U test between SNP groups identified by FarmCPU, BayesCπ, both, and neither. Number of causal SNPs is 256 and heritability is 0.5.

**Figure S1.**
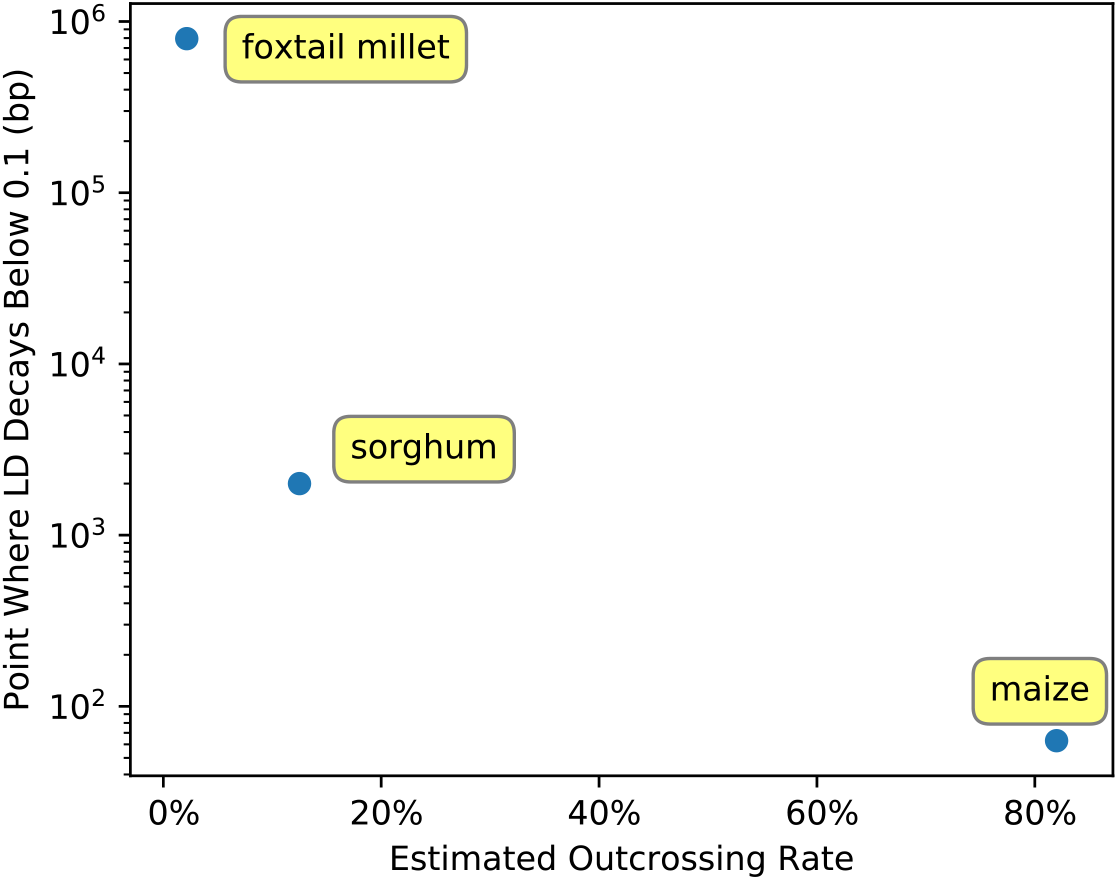
Relationship between LD decay and outcrossing rates reported from the literature for maize, sorghum, and foxtail millet. Estimated outcrossing rates for each species are taken from [35, 36, 37, 38, 39].

**Figure S2.**
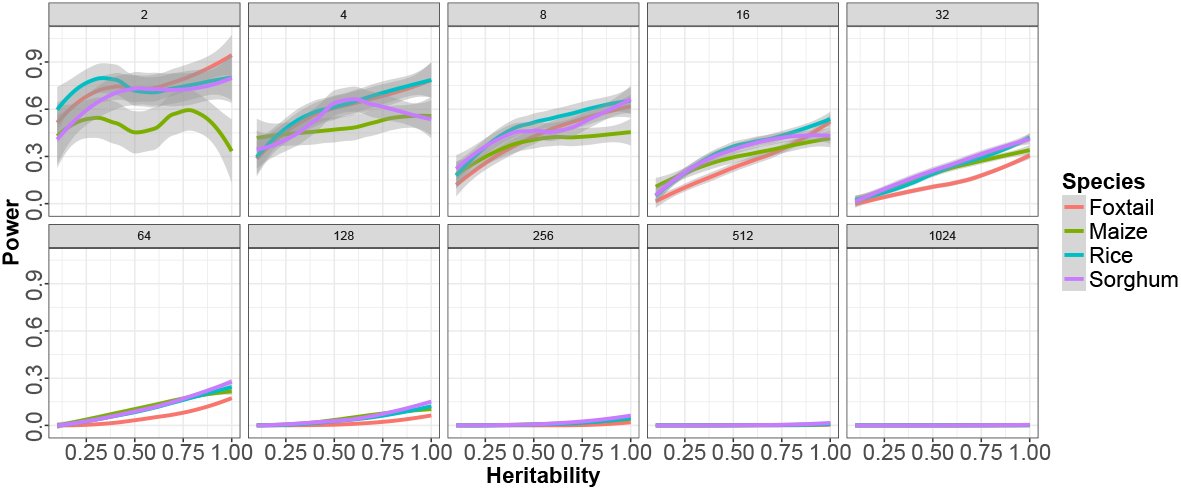
Relationship between the proportion of causal variants identified and heritability for traits controlled by different numbers of causal variants (2-1024 causal variants) in each species in an MLM-based GWAS. Positive calls were defined as those above a Bonferroni corrected p-value cutoff of 0.05.

**Figure S3.**
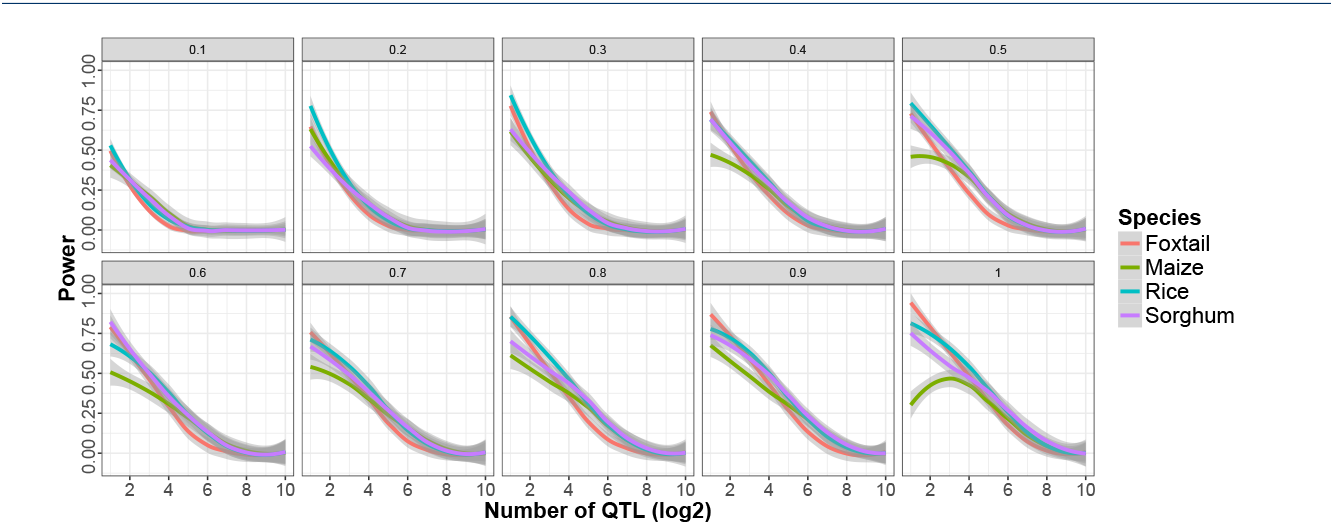
Relationship between the proportion of causal variants identified and the number of causal variants controlling a trait given different levels of heritability (0.1-1) in each species in an MLM-based GWAS. Positive calls were defined as those above a Bonferroni corrected p-value cutoff of 0.05.

**Figure S4.**
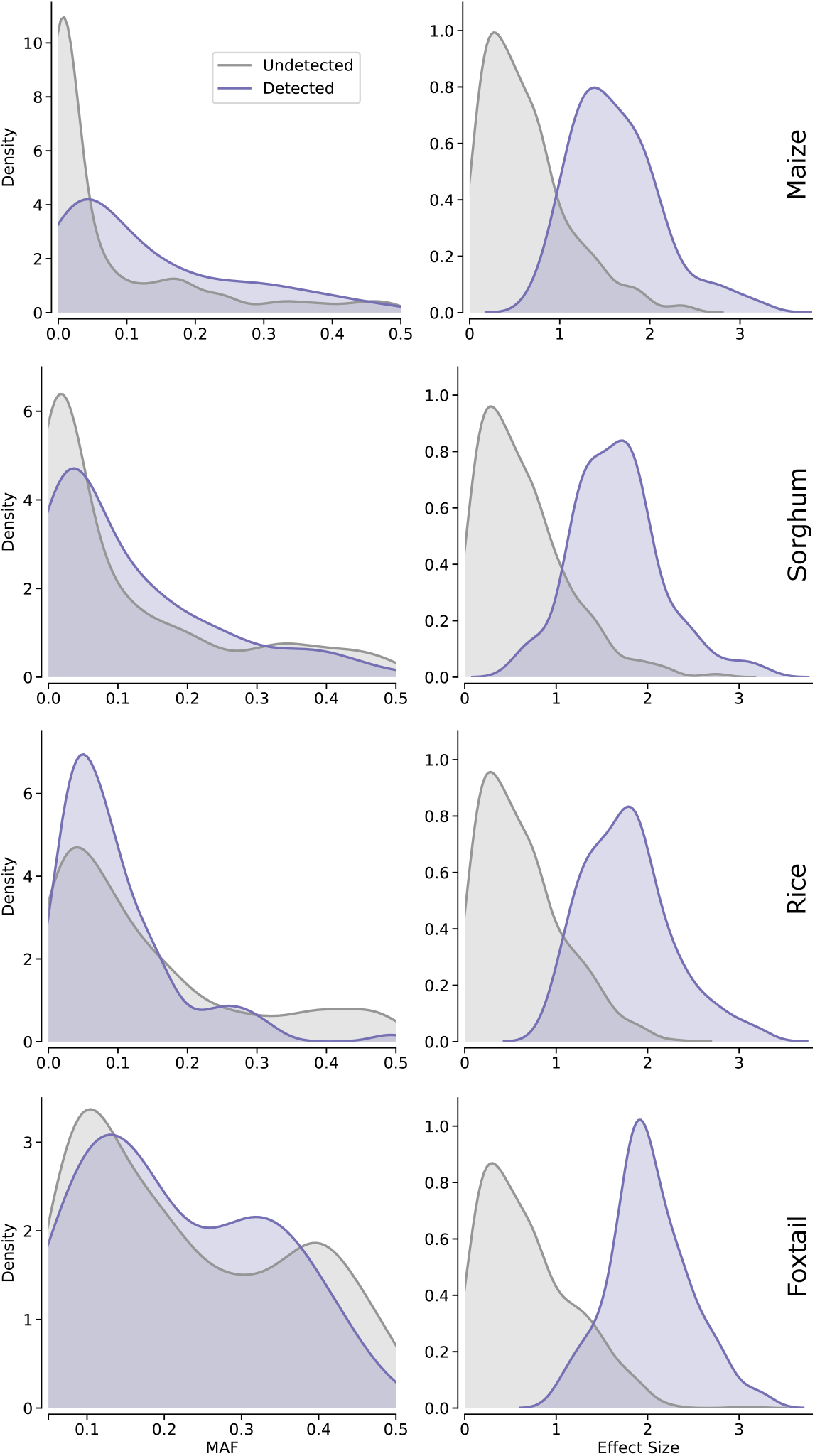
Distribution of minor allele frequency and effect size for true positive and false negative causal variants in each species. Data shown are from simulation here heritability is 0.7 and the number of simulated causal variants is 64. Positive calls were defined as those above a Bonferroni corrected p-value cutoff of 0.05.

**Figure S5.**
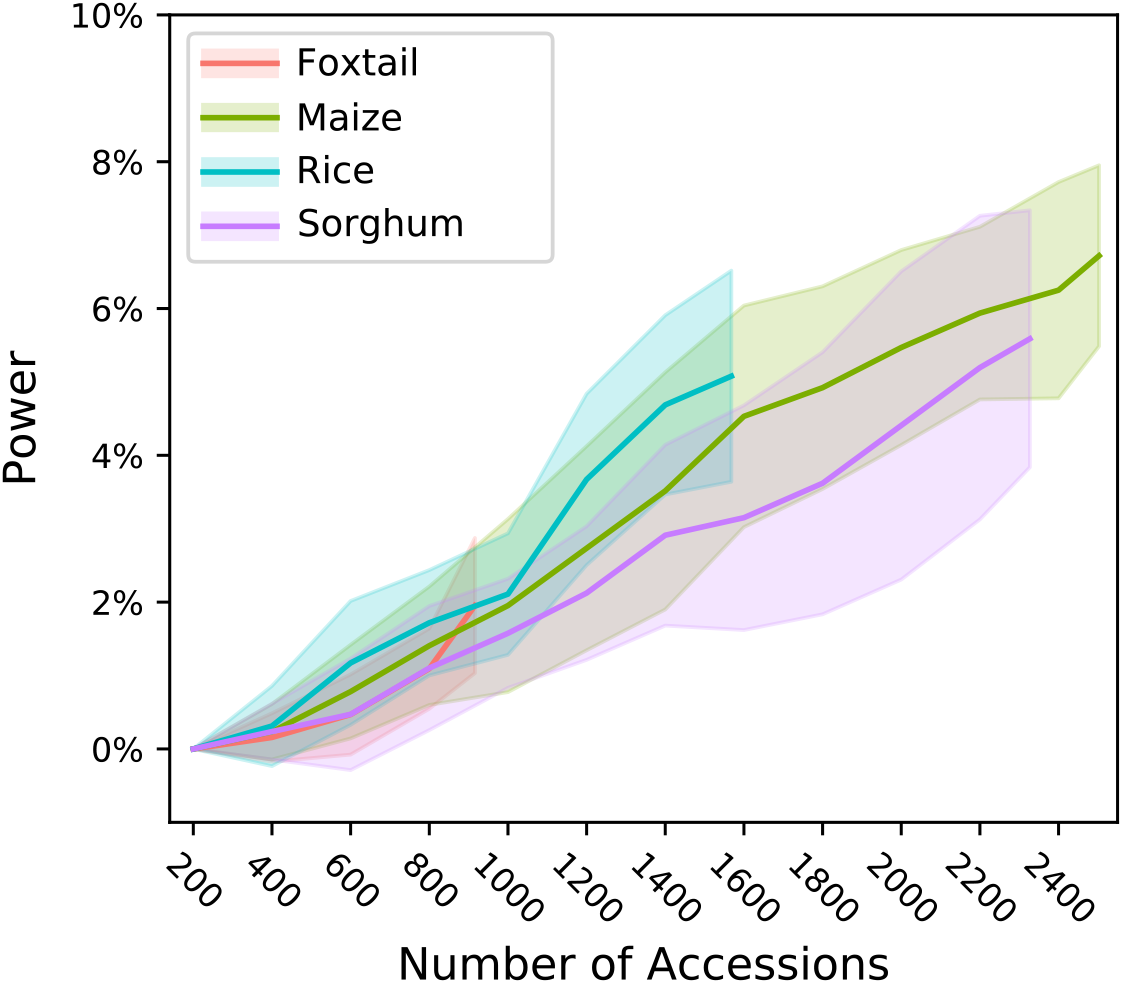
Proportion of causal variants identified by MLM based analysis at different population sizes in each species with 128 causal variants per simulation and heritability of 0.7. Positive calls were defined as those above a Bonferroni corrected p-value cutoff of 0.05.

**Figure S6.**
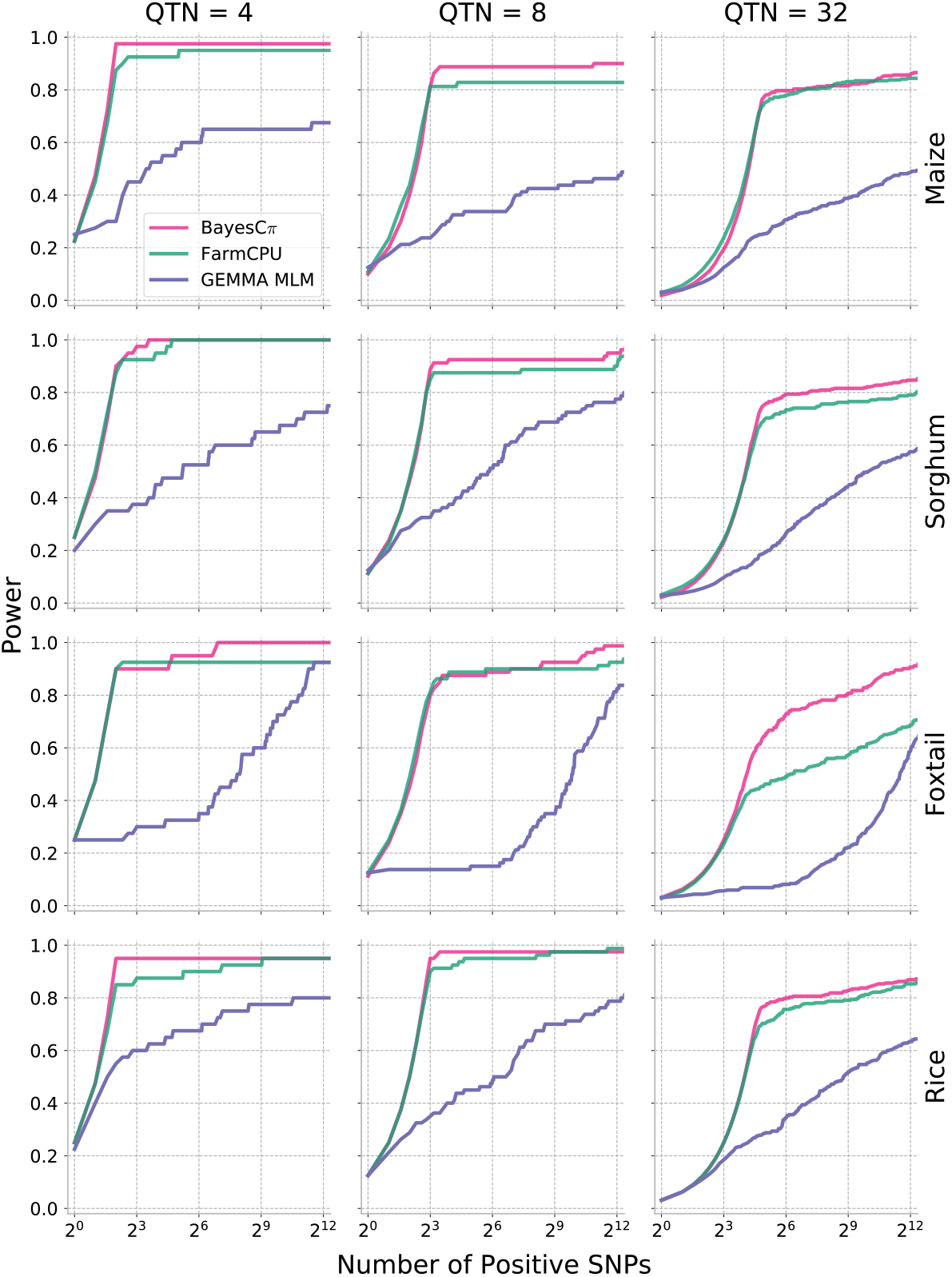
Relationship between the proportion of causal variants identified and the number of associated SNPs selected for MLM, MLMM, and Bayesian analysis for 4, 8, and 32 causal variants. Data shown are from simulations where trait heritability is 0.9.

**Figure S7.**
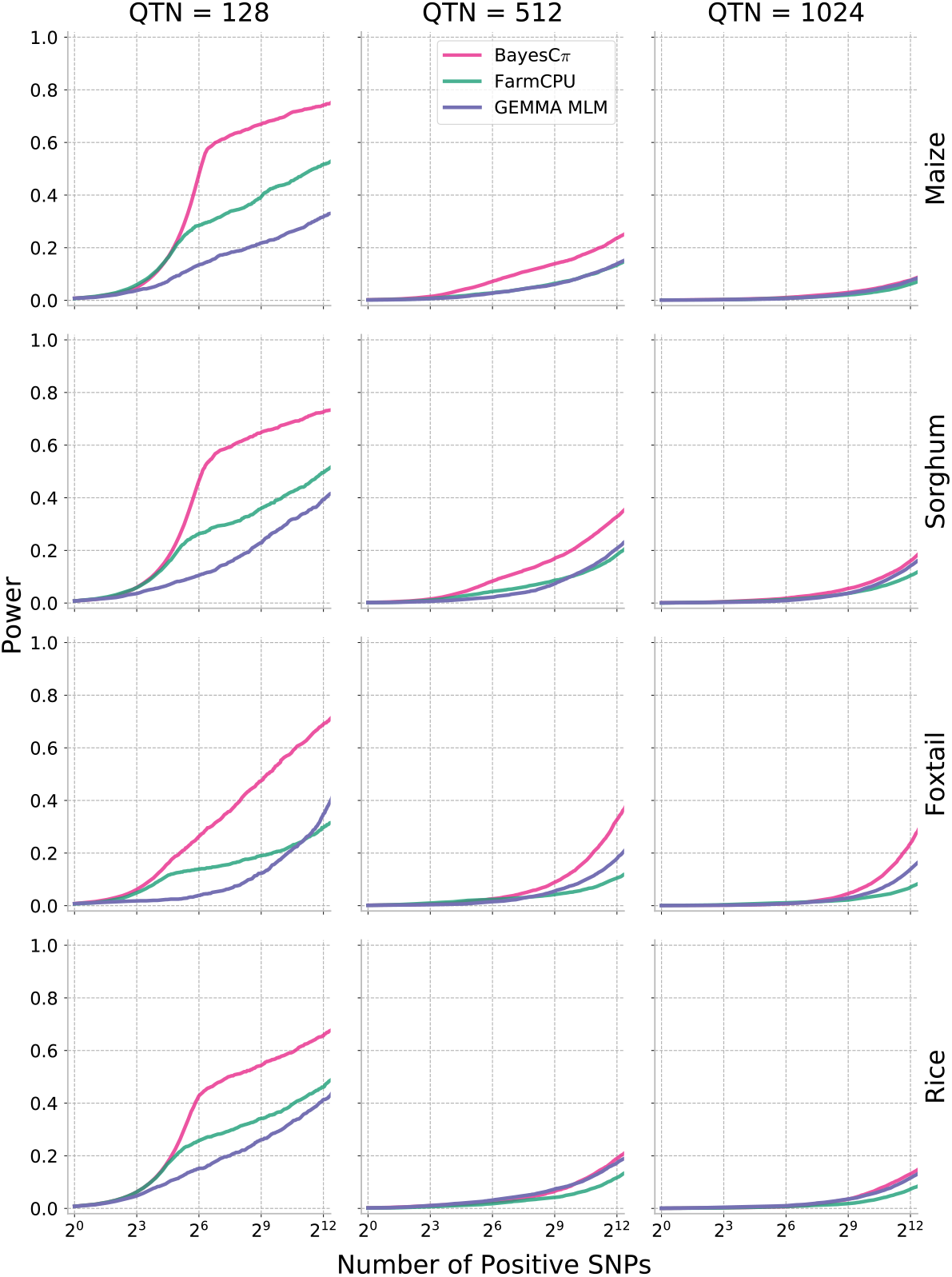
Relationship between the proportion of causal variants identified and the number of associated SNPs selected for MLM, MLMM, and Bayesian analysis for 128, 512, and 1024 causal variants. Data shown are from simulations where trait heritability is 0.9.

**Figure S8.**
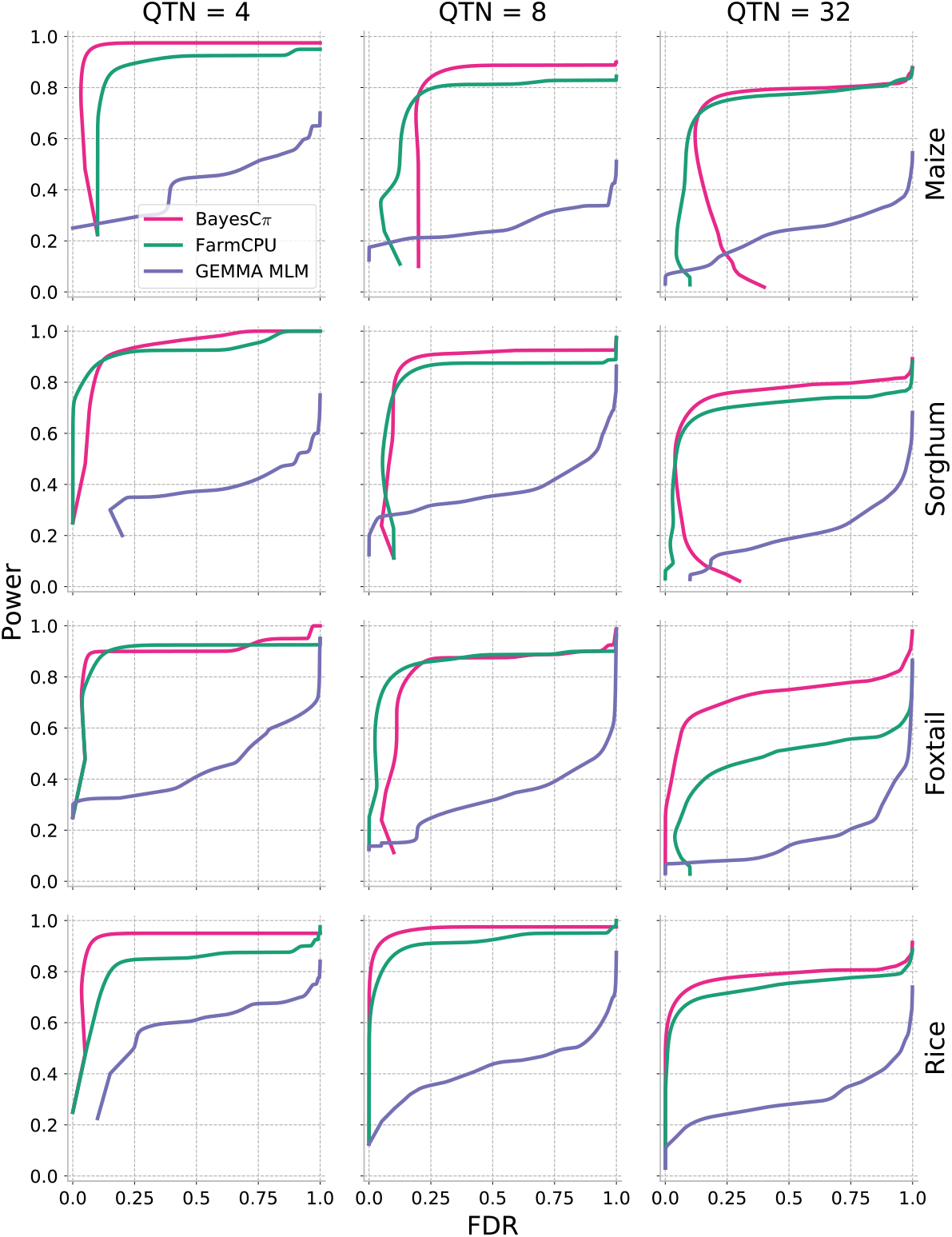
Relationship between false discovery rate and the number of associated SNPs selected for MLM, MLMM, and Bayesian analysis for 4, 8, and 32 causal variants. Data shown are from simulations where trait heritability is 0.9.

**Figure S9.**
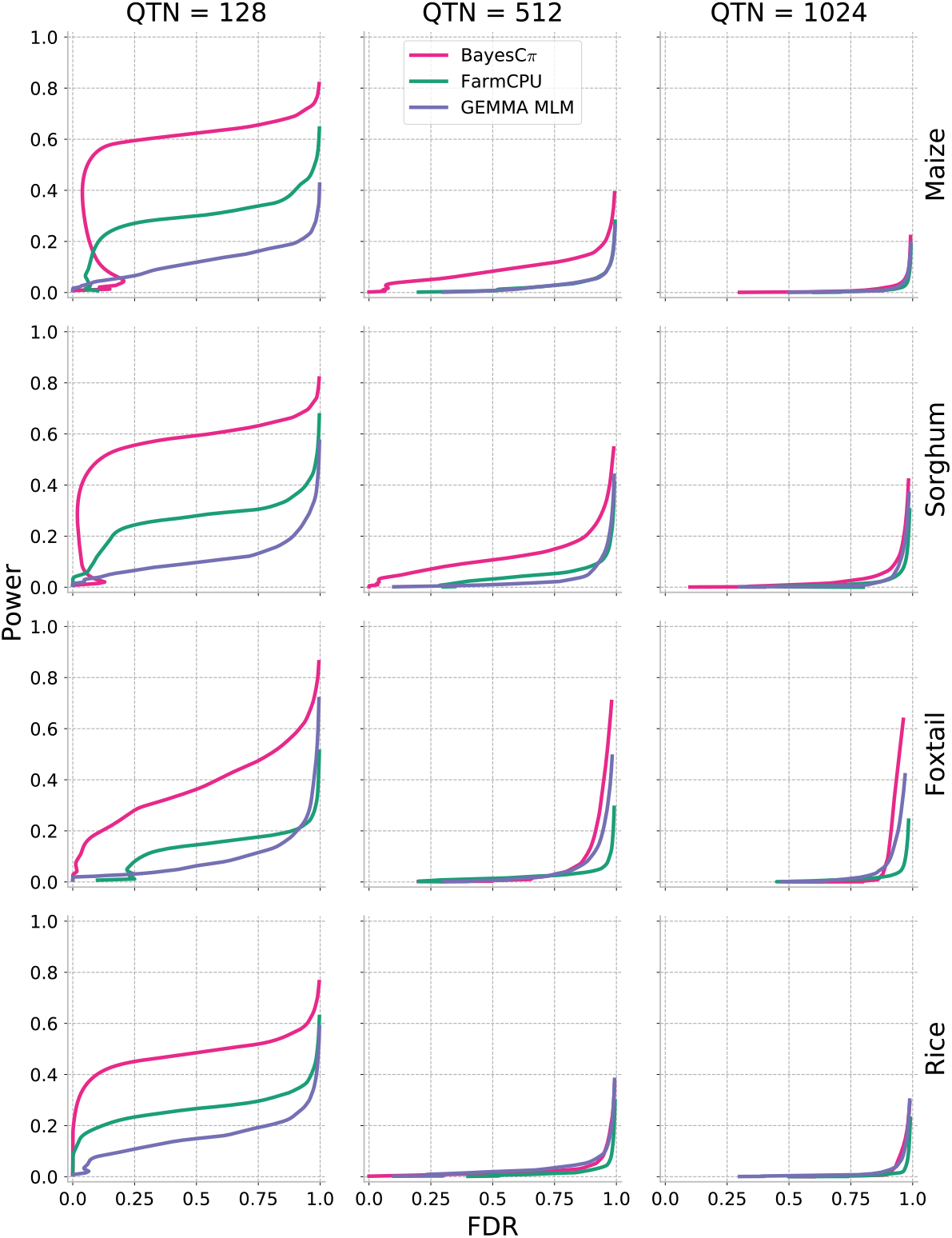
Relationship between false discovery rate and the number of associated SNPs selected for MLM, MLMM, and Bayesian analysis for 128, 512, and 1024 causal variants. Data shown are from simulations where trait heritability is 0.9.

**Figure S10.**
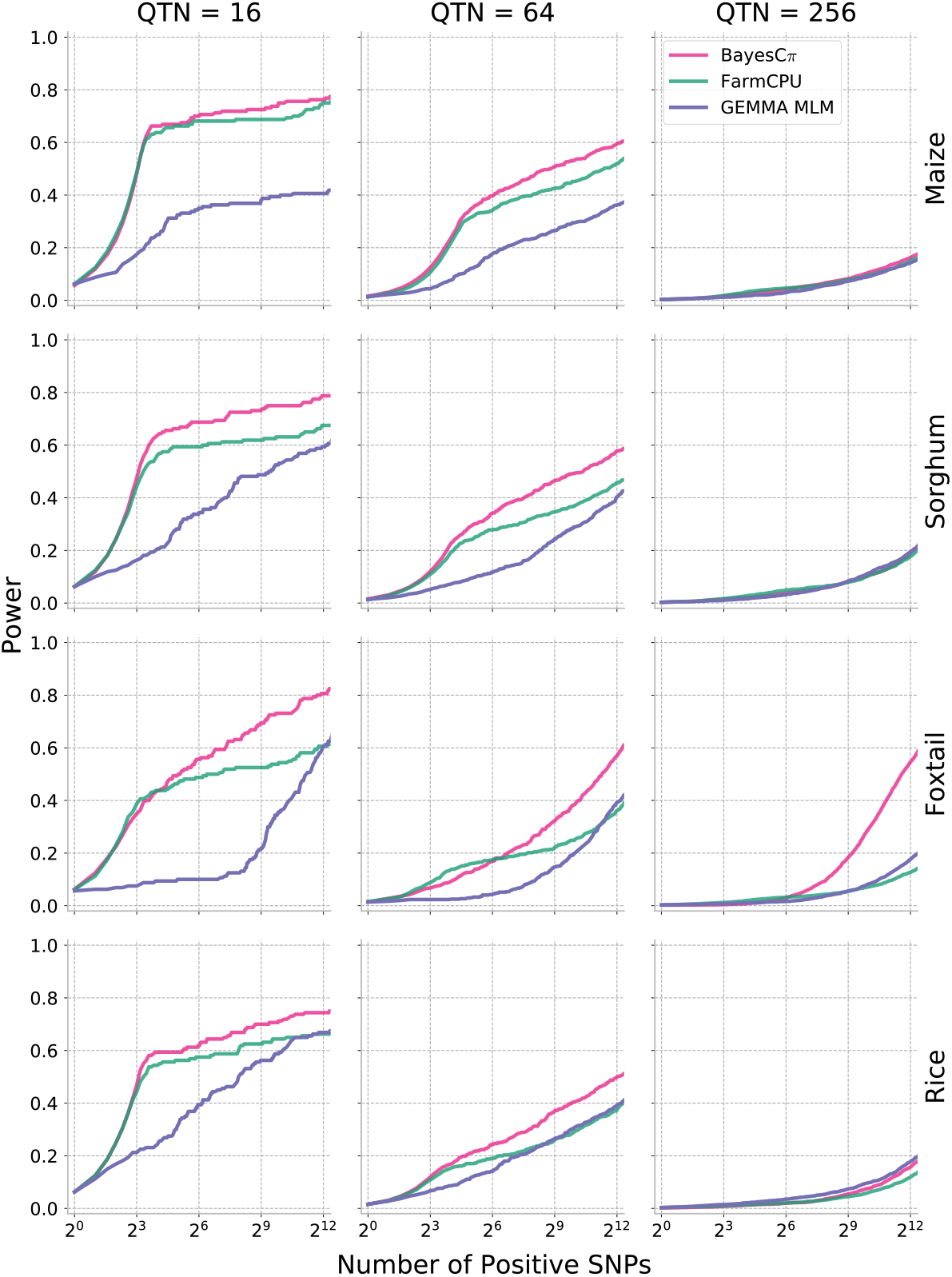
Relationship between the proportion of causal variants identified and the number of associated SNPs selected for MLM, MLMM, and Bayesian analysis for 16, 64, and 256 causal variants. Data shown are from simulations where trait heritability is 0.5.

**Figure S11.**
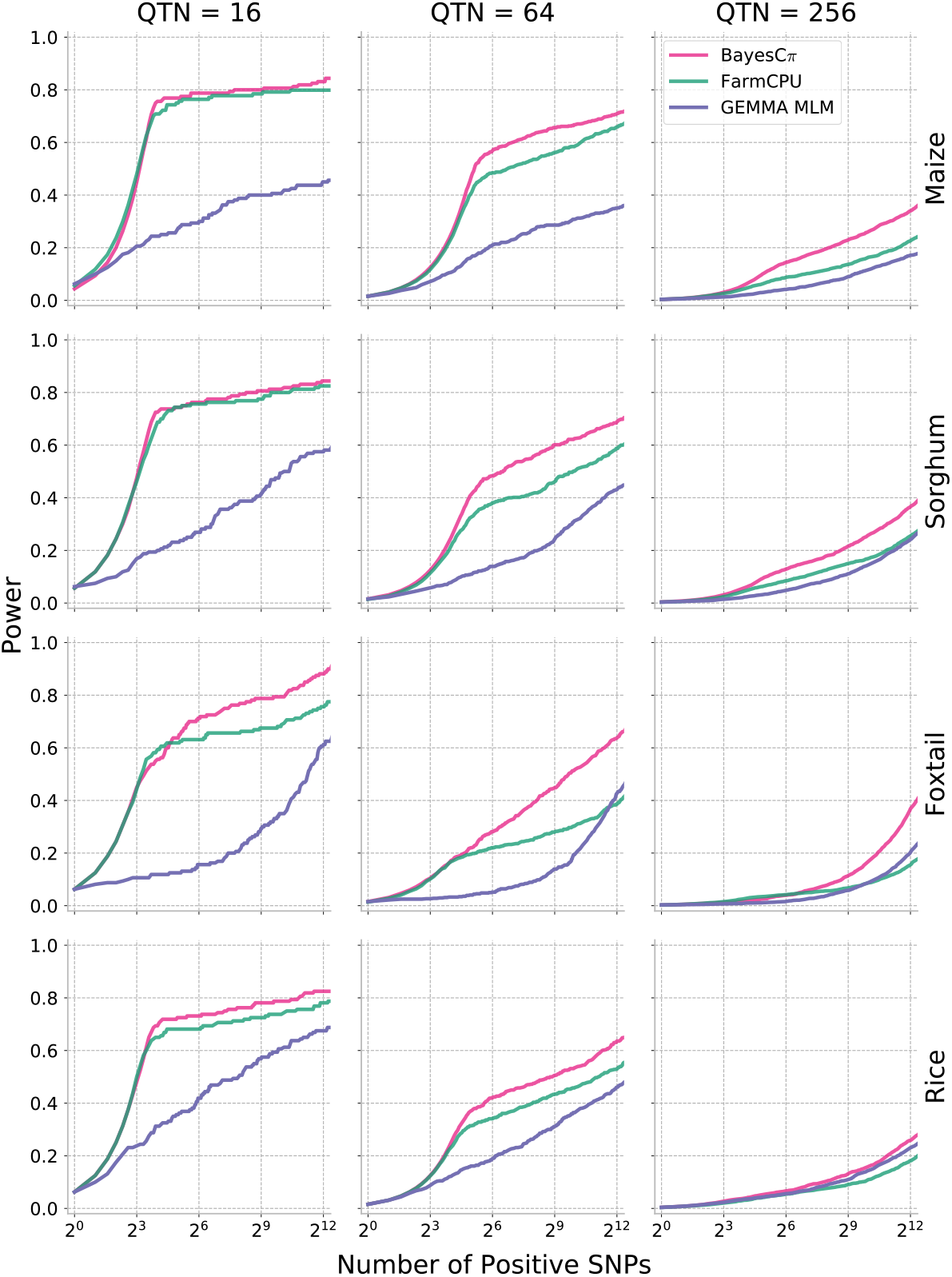
Relationship between the proportion of causal variants identified and the number of associated SNPs selected for MLM, MLMM, and Bayesian analysis for 16, 64, and 256 causal variants. Data shown are from simulations where trait heritability is 0.7.

**Figure S12.**
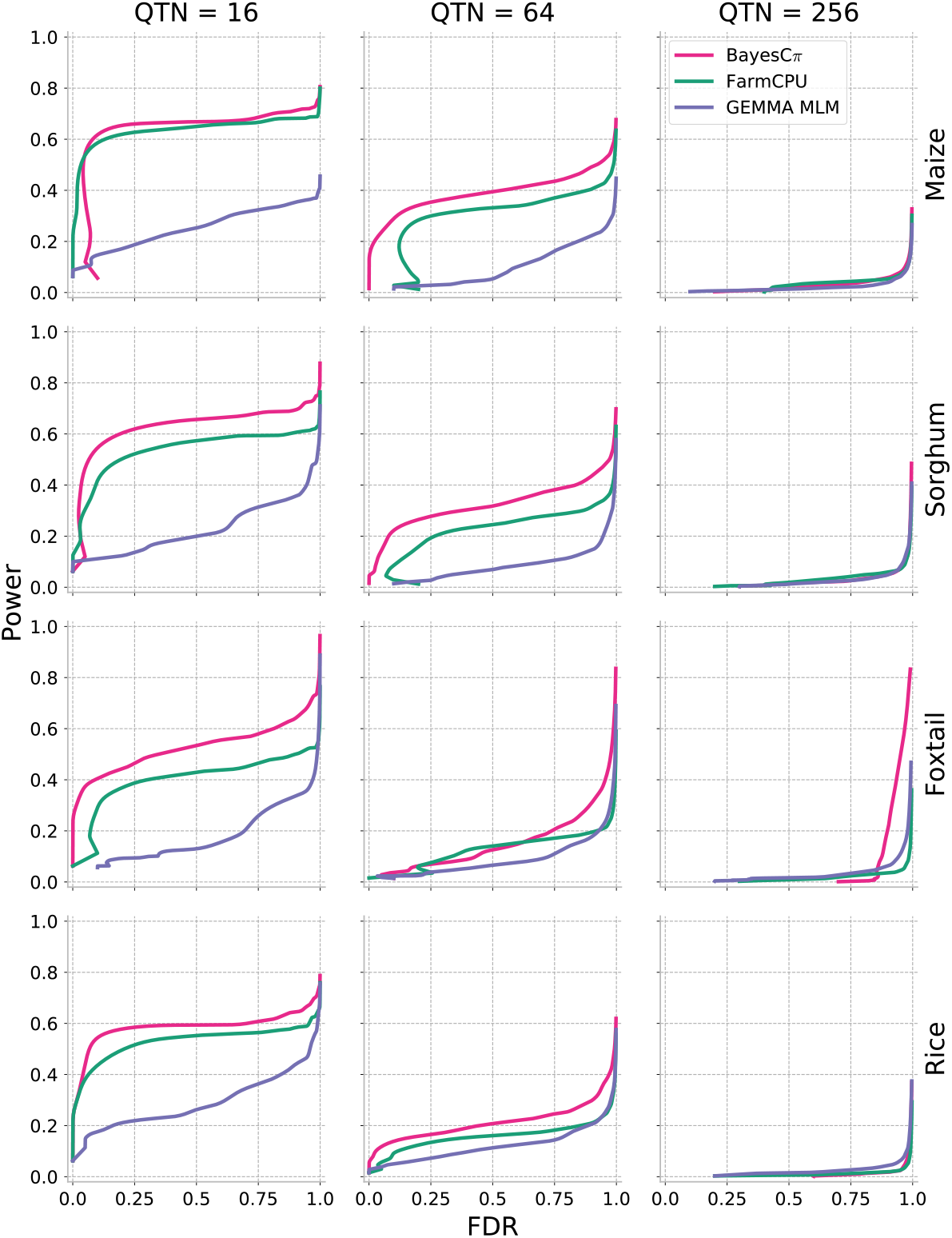
Relationship between false discovery rate and the number of associated SNPs selected for MLM, MLMM, and Bayesian analysis for 16, 64, and 256 causal variants. Data shown are from simulations where trait heritability is 0.5.

**Figure S13.**
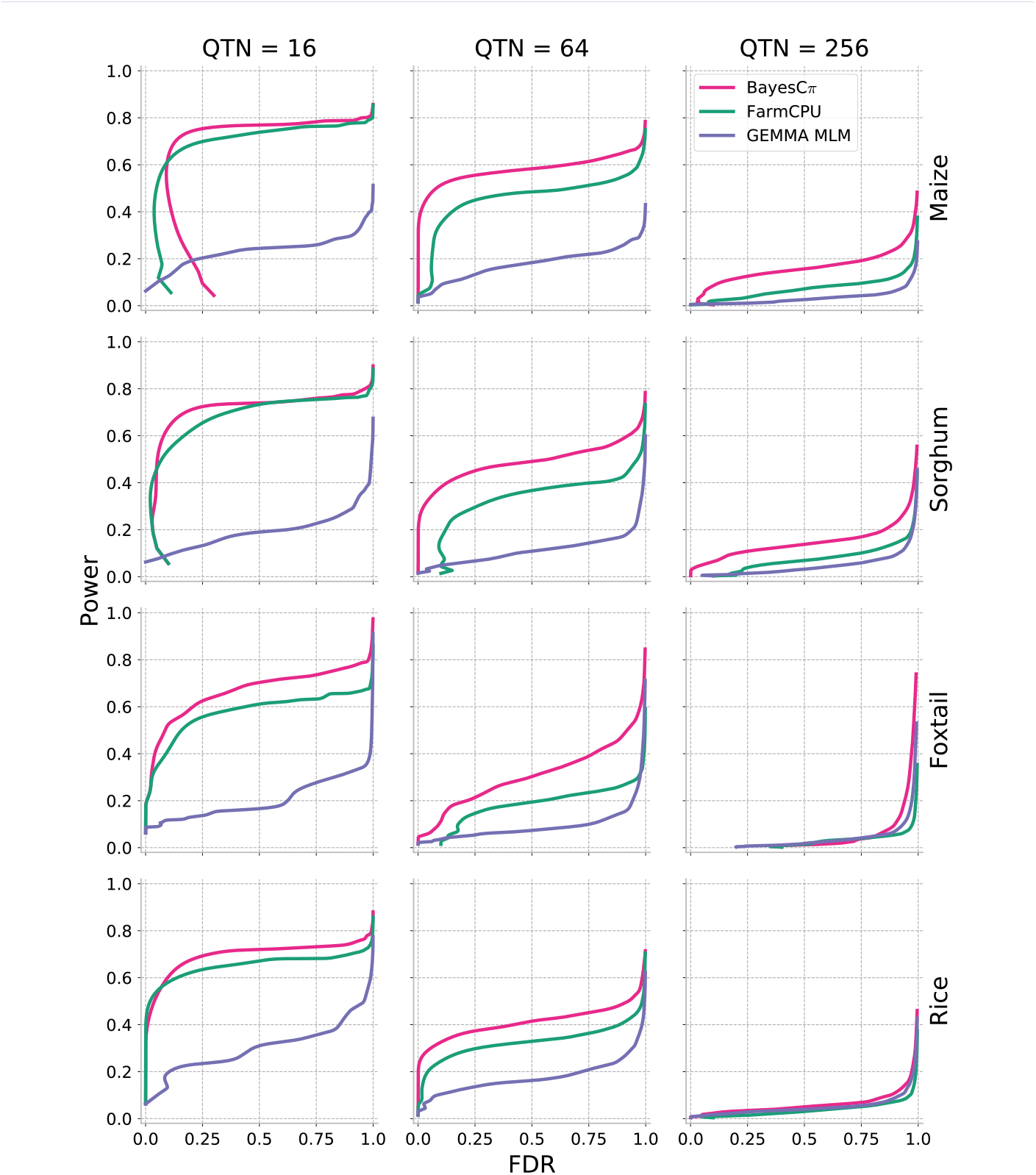
Relationship between false discovery rate and the number of associated SNPs selected for MLM, MLMM, and Bayesian analysis for 16, 64, and 256 causal variants. Data shown are from simulations where trait heritability is 0.7.

**Figure S14.**
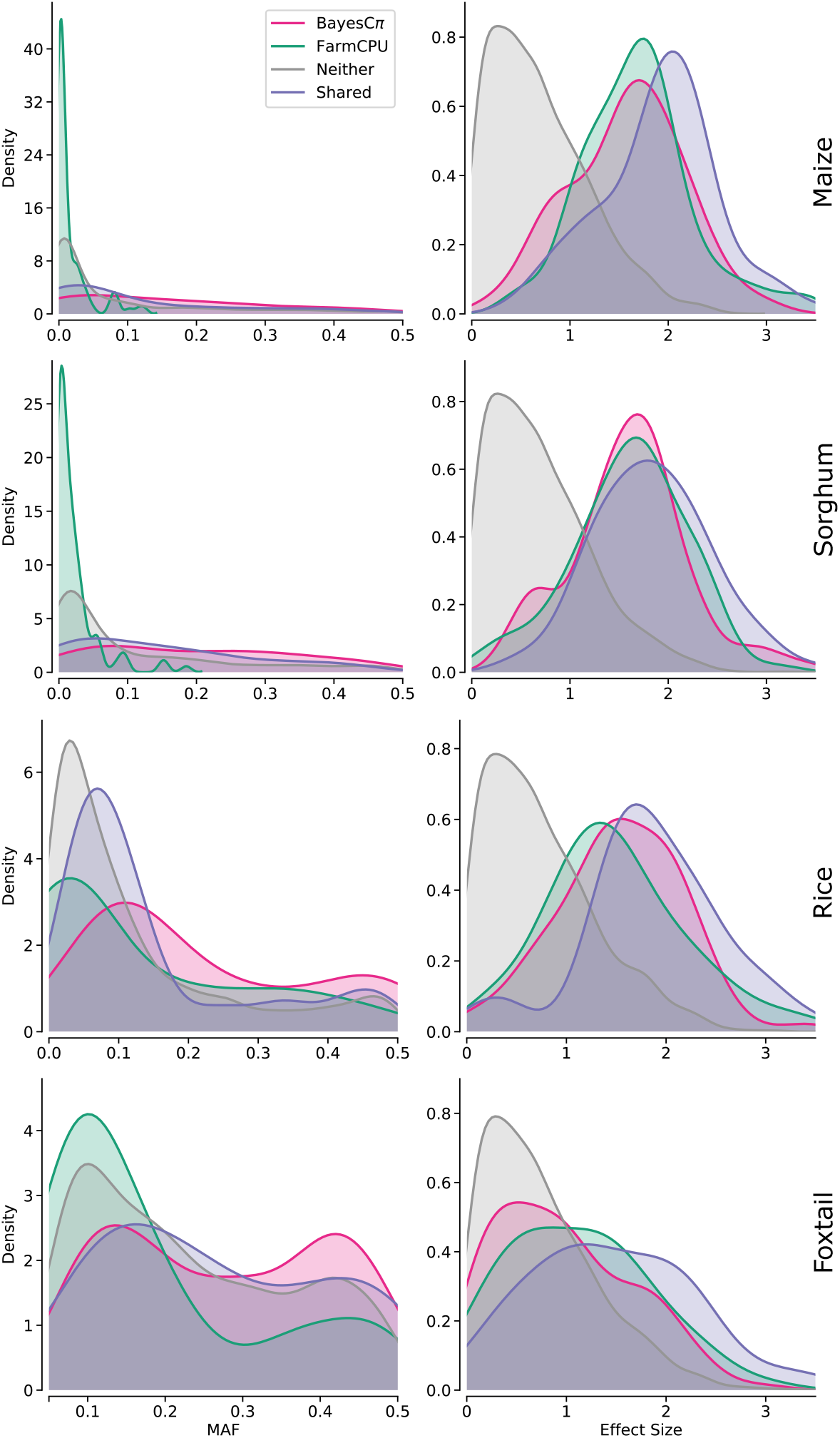
Differences in the characteristics of causal SNPs identified using either BayesCπ or FarmCPU for all populations in simulations with 256 causal variants and heritability of 0.5.

**Figure S15.**
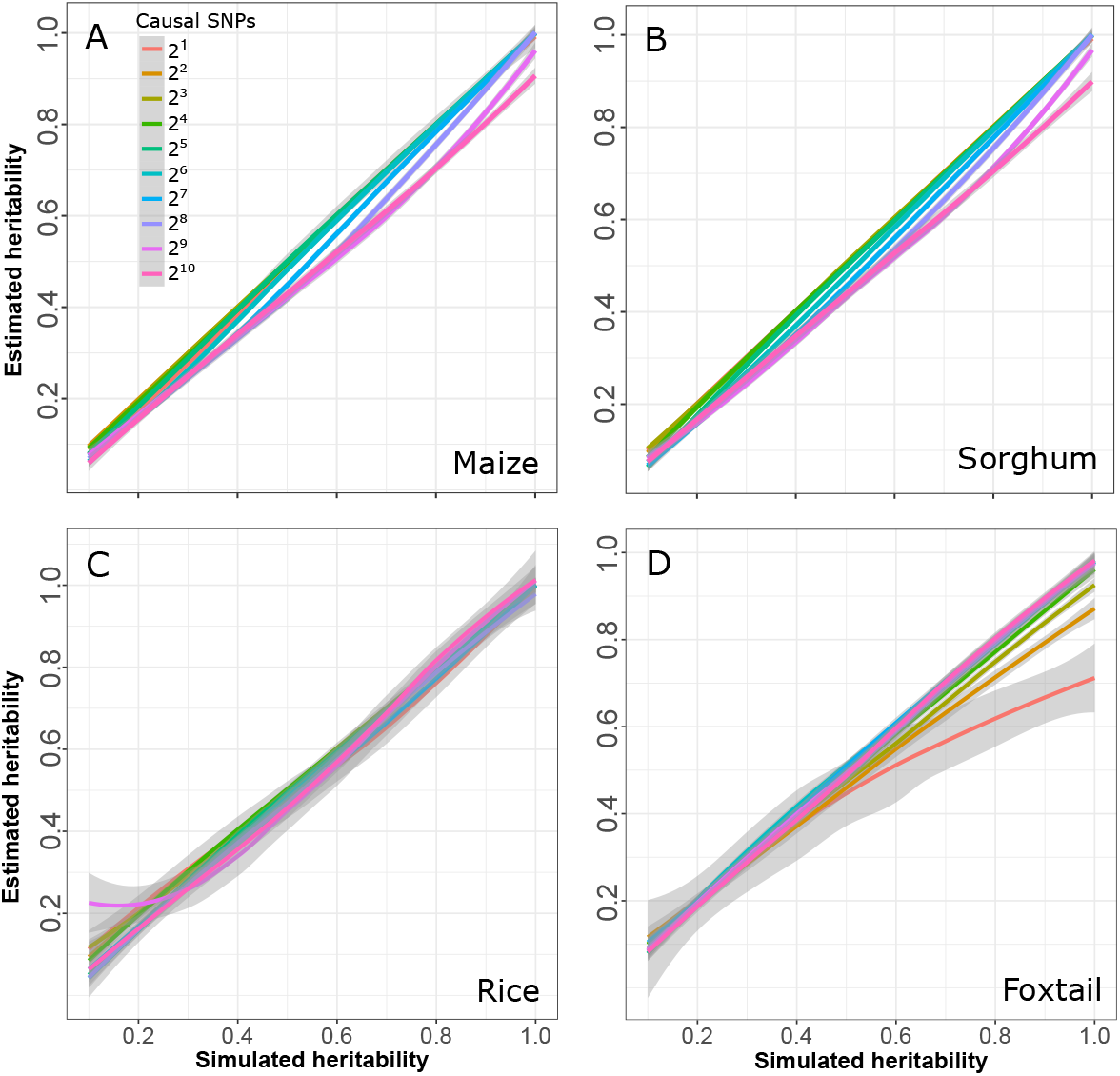
Relationship between simulated heritability and heritability estimates generated by BayesCπ for traits controlled by different numbers of causal variants. Grey lines indicate the 95% confidence intervals.

**Figure S16.**
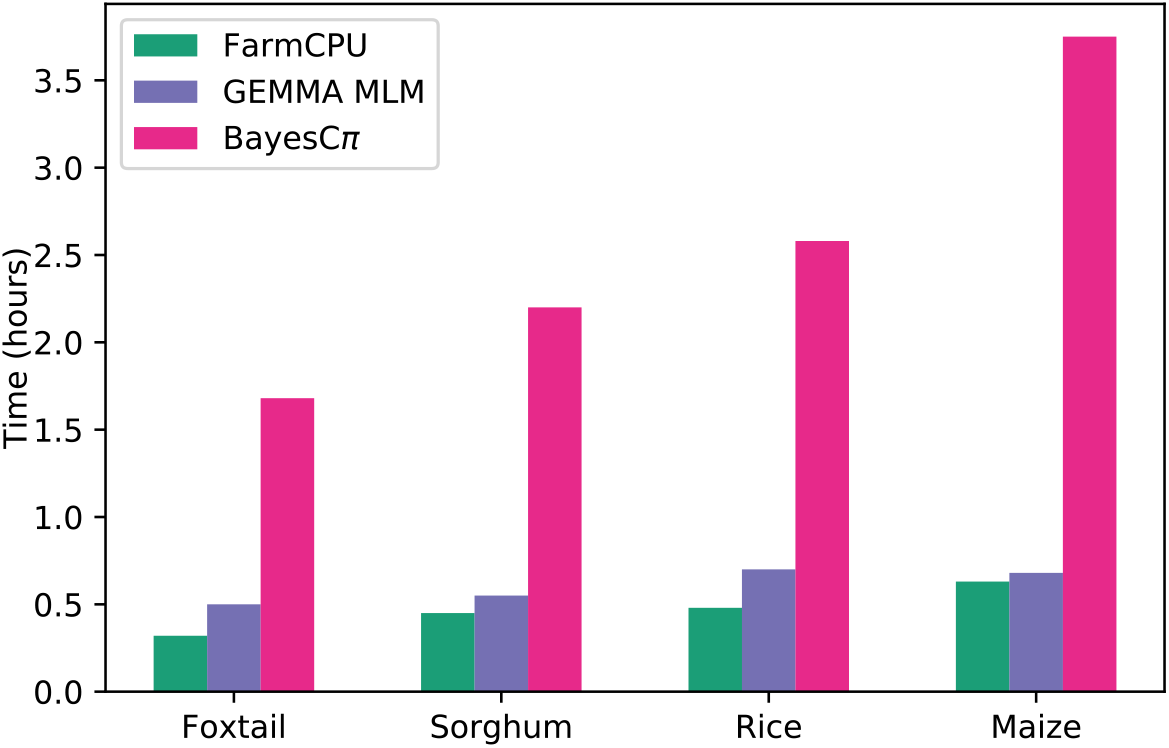
Average run time of a single GWAS analysis using each of the three methods evaluation in each of the four populations tested.

**Figure S17.**
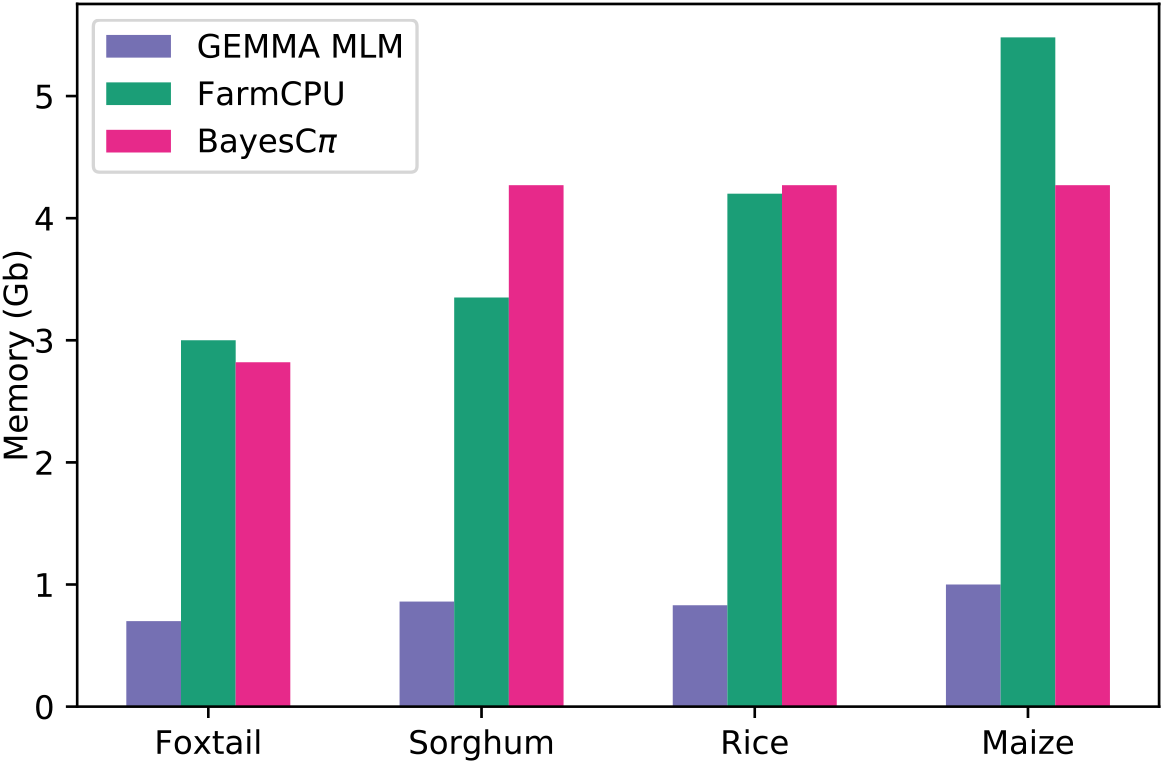
Average maximum memory use of a single GWAS analysis using each of the three methods evaluation in each of the four populations tested.

**Figure S18.**
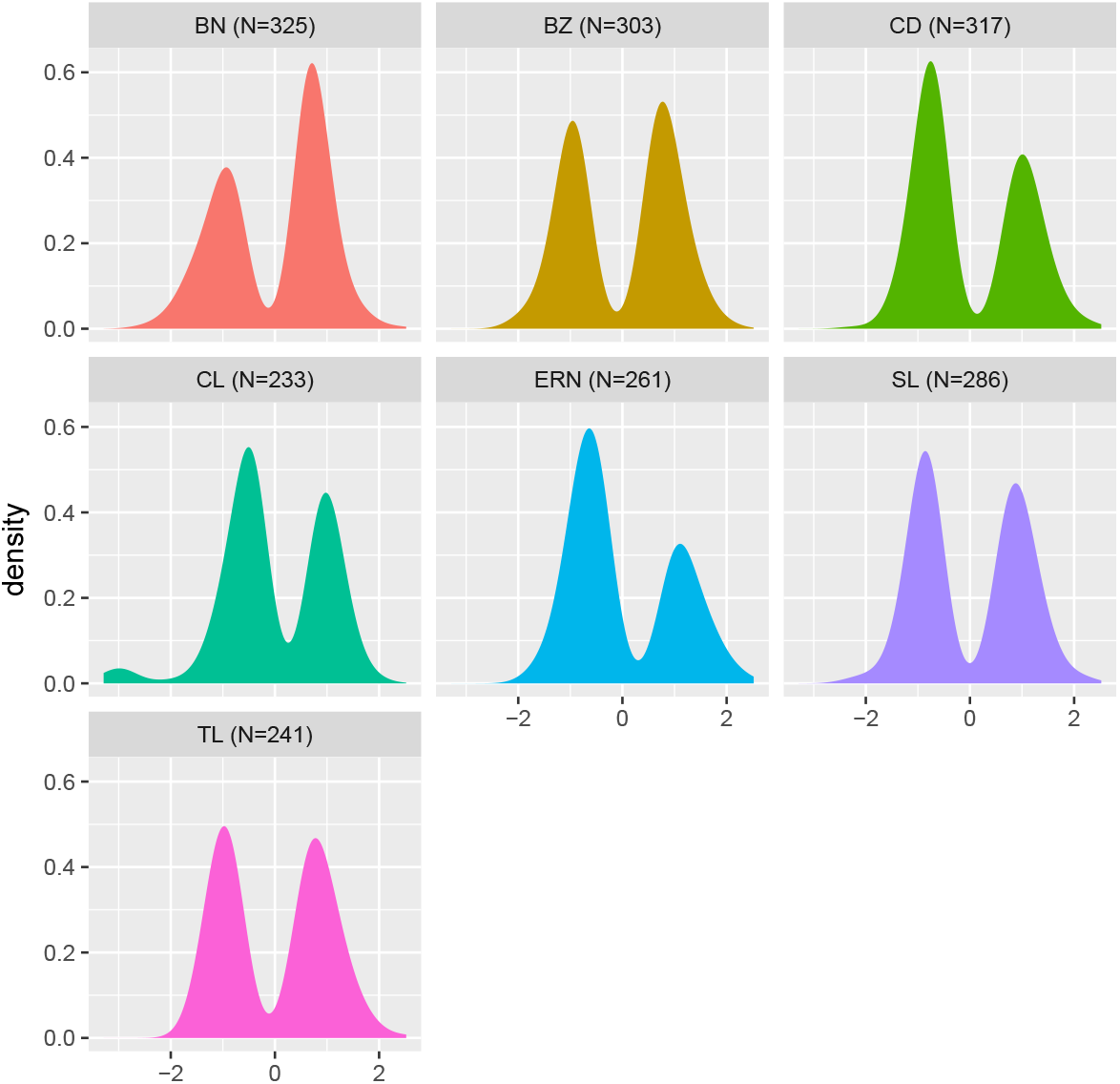
Empirically determined effect sizes for loci control seven different traits in maize. The SNP effects were obtained from [43]. In each case effects were normalized to a mean of 0 and a standard deviation of 1. In most cases, the distribution of effect sizes approximates the two tails of a normal distribution, with a missing center of unidentified small effect value SNPs. BN, branch number; BZ, branch zone; CD, cob diameter; CL, cob length; ERN, ear row number; SL, spike length; TL tassel length.

## References

1. Burton, P.R., Clayton, D.G., Cardon, L.R., Craddock, N., Deloukas, P., Duncanson, A., Kwiatkowski, D.P., McCarthy, M.I., Ouwehand, W.H., Samani, N.J., et al.: Genome-wide association study of 14,000 cases of seven common diseases and 3,000 shared controls. Nature 447(7145), 661–678 (2007)

2. Chen, W., Wang, W., Peng, M., Gong, L., Gao, Y., Wan, J., Wang, S., Shi, L., Zhou, B., Li, Z., et al.: Comparative and parallel genome-wide association studies for metabolic and agronomic traits in cereals. Nature communications 7 (2016)

3. Navarro, J.A.R., Willcox, M., Burgueño, J., Romay, C., Swarts, K., Trachsel, S., Preciado, E., Terron, A., Delgado, H.V., Vidal, V., et al.: A study of allelic diversity underlying flowering-time adaptation in maize landraces. Nature genetics 49(3), 476–480 (2017)

4. Jia, G., Huang, X., Zhi, H., Zhao, Y., Zhao, Q., Li, W., Chai, Y., Yang, L., Liu, K., Lu, H., et al.: A haplotype map of genomic variations and genome-wide association studies of agronomic traits in foxtail millet (setaria italica). Nature genetics 45(8), 957–961 (2013)

5. Lasky, J.R., Upadhyaya, H.D., Ramu, P., Deshpande, S., Hash, C.T., Bonnette, J., Juenger, T.E., Hyma, K., Acharya, C., Mitchell, S.E., et al.: Genome-environment associations in sorghum landraces predict adaptive traits. Science advances 1(6), 1400218 (2015)

6. Maher, B.: Personal genomes: The case of the missing heritability. Nature News 456(7218), 18–21 (2008)

7. Manolio, T.A., Collins, F.S., Cox, N.J., Goldstein, D.B., Hindorff, L.A., Hunter, D.J., McCarthy, M.I., Ramos, E.M., Cardon, L.R., Chakravarti, A., et al.: Finding the missing heritability of complex diseases. Nature 461(7265), 747–753 (2009)

8. Visscher, P.M., Yang, J., Goddard, M.E.: A commentary on ‘common snps explain a large proportion of the heritability for human height’by yang et al. (2010). Twin Research and Human Genetics 13(6), 517–524 (2010)

9. Gerasimova, A., Chavez, L., Li, B., Seumois, G., Greenbaum, J., Rao, A., Vijayanand, P., Peters, B.: Predicting cell types and genetic variations contributing to disease by combining gwas and epigenetic data. PloS one 8(1), 54359 (2013)

10. Visscher, P.M., Hill, W.G., Wray, F.R.: Heritability in the genomics era—concepts and misconceptions. Nature reviews genetics 9(4), 255 (2008)

11. Moellers, T.C., Singh, A., Zhang, J., Brungardt, J., Kabbage, M., Mueller, D.S., Grau, C.R., Ranjan, A., Smith, D.L., Chowda-Reddy, R., et al.: Main and epistatic loci studies in soybean for sclerotinia sclerotiorum resistance reveal multiple modes of resistance in multi-environments. Scientific reports 7(1), 3554 (2017)

12. Zhang, J., Singh, A., Mueller, D.S., Singh, A.K.: Genome-wide association and epistasis studies unravel the genetic architecture of sudden death syndrome resistance in soybean. The Plant Journal 84(6), 1124–1136 (2015)

13. McCarroll, S.A.: Extending genome-wide association studies to copy-number variation. Human molecular genetics 17(R2), 135–142 (2008)

14. Jakobsdottir, J., Gorin, M.B., Conley, Y.P., Ferrell, R.E., Weeks, D.E.: Interpretation of genetic association studies: markers with replicated highly significant odds ratios may be poor classifiers. PLoS genetics 5(2), 1000337 (2009)

15. Pritchard, J.K.: Are rare variants responsible for susceptibility to complex diseases? The American Journal of Human Genetics 69(1), 124–137 (2001)

16. Huang, X., Zhao, Y., Li, C., Wang, A., Zhao, Q., Li, W., Guo, Y., Deng, L., Zhu, C., Fan, D., et al.: Genome-wide association study of flowering time and grain yield traits in a worldwide collection of rice germplasm. Nature genetics 44(1), 32–39 (2012)

17. Boyle, E.A., Li, Y.I., Pritchard, J.K.: An expanded view of complex traits: from polygenic to omnigenic. Cell 169(7), 1177–1186 (2017)

18. Kang, H.M., Sul, J.H., Zaitlen, F.A., Kong, S.-y., Freimer, F.B., Sabatti, C., Eskin, E., et al.: Variance component model to account for sample structure in genome-wide association studies. Nature genetics 42(4), 348–354 (2010)

19. Zhang, Z., Ersoz, E., Lai, C.-Q., Todhunter, R.J., Tiwari, H.K., Gore, M.A., Bradbury, P.J., Yu, J., Arnett, D.K., Ordovas, J.M., et al.: Mixed linear model approach adapted for genome-wide association studies. Nature genetics 42(4), 355–360 (2010)

20. Lippert, C., Listgarten, J., Liu, Y., Kadie, C.M., Davidson, R.I., Heckerman, D.: Fast linear mixed models for genome-wide association studies. Nature methods 8(10), 833–835 (2011)

21. Zhou, X., Stephens, M.: Genome-wide efficient mixed-model analysis for association studies. Nature genetics 44(7), 821–824 (2012)

22. Segura, V., Vilhj’lmsson, B.J., Platt, A., Korte, A., Seren, Ü., Long, Q., Nordborg, M.: An efficient multi-locus mixed-model approach for genome-wide association studies in structured populations. Nature genetics 44(7), 825–830 (2012)

23. Liu, X., Huang, M., Fan, B., Buckler, E.S., Zhang, Z.: Iterative usage of fixed and random effect models for powerful and efficient genome-wide association studies. PLoS genetics 12(2), 1005767 (2016)

24. Schnable, P.S., Kusmec, A.: Farmcpupp: Efficient large-scale gwas. bioRxiv, 238832 (2017)

25. Hayes, B., Goddard, M., et al.: Prediction of total genetic value using genome-wide dense marker maps. Genetics 157(4), 1819–1829 (2001)

26. Bernardo, R., Yu, J.: Prospects for genomewide selection for quantitative traits in maize. Crop Science 47(3), 1082–1090 (2007)

27. Piepho, H.-P.: Ridge regression and extensions for genomewide selection in maize. Crop Science 49(4), 1165–1176 (2009)

28. Verbyla, K.L., Hayes, B.J., Bowman, P.J., Goddard, M.E.: Accuracy of genomic selection using stochastic search variable selection in australian holstein friesian dairy cattle. Genetics research 91(5), 307–311 (2009)

29. Rohan L. Fernando, D.G.: Bayesian methods applied to gwas. In: Cedric Gondro, J.v.d.W. Ben Hayes (ed.) Genome-Wide Association Studies and Genomic Prediction vol. 1019, pp. 237–274. Springer, ??? (2013)

30. Sun, X., Habier, D., Fernando, R.L., Garrick, D.J., Dekkers, J.C.: Genomic breeding value prediction and qtl mapping of qtlmas2010 data using bayesian methods. In: BMC Proceedings, vol. 5, p. 13 (2011). BioMed Central

31. Fan, B., Onteru, S.K., Du, Z.-Q., Garrick, D.J., Stalder, K.J., Rothschild, M.F.: Genome-wide association study identifies loci for body composition and structural soundness traits in pigs. PloS one 6(2), 14726 (2011)

32. Peters, S., Kizilkaya, K., Garrick, D., Fernando, R., Reecy, J., Weaber, R., Silver, G., Thomas, M.: Bayesian genome-wide association analysis of growth and yearling ultrasound measures of carcass traits in brangus heifers. Journal of animal science 90(10), 3398–3409 (2012)

33. Romay, M.C., Millard, M.J., Glaubitz, J.C., Peiffer, J.A., Swarts, K.L., Casstevens, T.M., Elshire, R.J., Acharya, C.B., Mitchell, S.E., Flint-Garcia, S.A., et al.: Comprehensive genotyping of the usa national maize inbred seed bank. Genome biology 14(6), 55 (2013)

34. McCouch, S.R., Wright, M.H., Tung, C.-W., Maron, L.G., McNally, K.L., Fitzgerald, M., Singh, N., DeClerck, G., Agosto-Perez, F., Korniliev, P., et al.: Open access resources for genome-wide association mapping in rice. Nature communications 7, 10532 (2016)

35. Gutierrez, M.G., Sprague, G.: Randomness of mating in isolated polycross plantings of maize. Genetics 44(6), 1075–1082 (1959)

36. Djè, Y., Heuertz, M., Ater, M., Lefèbvre, C., Vekemans, X.: In situ estimation of outcrossing rate in sorghum landraces using microsatellite markers. Euphytica 138(3), 205–212 (2004)

37. Barnaud, A., Trigueros, G., McKey, D., Joly, H.: High outcrossing rates in fields with mixed sorghum landraces: how are landraces maintained? Heredity 101(5), 445 (2008)

38. Wang, C., Chen, J., Zhi, H., Yang, L., Li, W., Wang, Y., Li, H., Zhao, B., Chen, M., Diao, X.: Population genetics of foxtail millet and its wild ancestor. BMC genetics 11(1), 90 (2010)

39. Hufford, M.B., Gepts, P., ROSS-IBARRA, J.: Influence of cryptic population structure on observed mating patterns in the wild progenitor of maize (zea mays ssp. parviglumis). Molecular ecology 20(1), 46–55 (2011)

40. Habier, D., Fernando, R.L., Kizilkaya, K., Garrick, D.J.: Extension of the bayesian alphabet for genomic selection. BMC bioinformatics 12(1), 186 (2011)

41. Bernardo, R.: Bandwagons i, too, have known. Theoretical and Applied Genetics 129(12), 2323–2332 (2016)

42. Buckler, E.S., Holland, J.B., Bradbury, P.J., Acharya, C.B., Brown, P.J., Browne, C., Ersoz, E., Flint-Garcia, S., Garcia, A., Glaubitz, J.C., et al.: The genetic architecture of maize flowering time. Science 325(5941), 714–718 (2009)

43. Brown, P.J., Upadyayula, N., Mahone, G.S., Tian, F., Bradbury, P.J., Myles, S., Holland, J.B., Flint-Garcia, S., McMullen, M.D., Buckler, E.S., et al.: Distinct genetic architectures for male and female inflorescence traits of maize. PLoS genetics 7(11), 1002383 (2011)

44. Kremling, K.A., Chen, S.-Y., Su, M.-H., Lepak, N.K., Romay, M.C., Swarts, K.L., Lu, F., Lorant, A., Bradbury, P.J., Buckler, E.S.: Dysregulation of expression correlates with rare-allele burden and fitness loss in maize. Nature (2018)

45. Fernando, R., Toosi, A., Wolc, A., Garrick, D., Dekkers, J.: Application of whole-genome prediction methods for genome-wide association studies: a bayesian approach. Journal of Agricultural, Biological and Environmental Statistics 22(2), 172–193 (2017)

46. Xia, F., Zhang, M.J., Zou, J.Y., Tse, D.: Neuralfdr: Learning discovery thresholds from hypothesis features. Ini Advances in Neural Information Processing Systems, pp. 1540–1549 (2017)

47. Bennetzen, J.L., Schmutz, J., Wang, H., Percifield, R., Hawkins, J., Pontaroli, A.C., Estep, M., Feng, L., Vaughn, J.N., Grimwood, J., et al.: Reference genome sequence of the model plant setaria. Nature biotechnology 30(6), 555–561 (2012)

48. Paterson, A.H., Bowers, J.E., Bruggmann, R., Dubchak, I., Grimwood, J., Gundlach, H., Haberer, G., Hellsten, U., Mitros, T., Poliakov, A., et al.: The sorghum bicolor genome and the diversification of grasses. Nature 457(7229), 551–556 (2009)

49. Schnable, P.S., Ware, D., Fulton, R.S., Stein, J.C., Wei, F., Pasternak, S., Liang, C., Zhang, J., Fulton, L., Graves, T.A., et al.: The b73 maize genome: complexity, diversity, and dynamics. science 326(5956), 1112–1115 (2009)

50. Browning, B.L., Browning, S.R.: Genotype imputation with millions of reference samples. The American Journal of Human Genetics 98(1), 116–126 (2016)

51. Bradbury, P.J., Zhang, Z., Kroon, D.E., Casstevens, T.M., Ramdoss, Y., Buckler, E.S.: Tassel: software for association mapping of complex traits in diverse samples. Bioinformatics 23(19), 2633–2635 (2007)

52. Purcell, S., Neale, B., Todd-Brown, K., Thomas, L., Ferreira, M.A., Bender, D., Maller, J., Sklar, P., De Bakker, P.I., Daly, M.J., et al.: Plink: a tool set for whole-genome association and population-based linkage analyses. The American Journal of Human Genetics 81(3), 559–575 (2007)

